# Analysis of donor pancreata defines the transcriptomic signature and microenvironment of early pre-neoplastic pancreatic lesions

**DOI:** 10.1101/2023.01.13.523300

**Authors:** Eileen S. Carpenter, Ahmed M. Elhossiny, Padma Kadiyala, Jay Li, Jake McGue, Brian Griffith, Yaqing Zhang, Jacob Edwards, Sarah Nelson, Fatima Lima, Katelyn L. Donahue, Wenting Du, Allison C. Bischoff, Danyah Alomari, Hannah Watkoske, Michael Mattea, Stephanie The, Carlos Espinoza, Meredith Barrett, Christopher J. Sonnenday, Nicholas Olden, Nicole Peterson, Valerie Gunchick, Vaibhav Sahai, Arvind Rao, Filip Bednar, Jiaqi Shi, Timothy L. Frankel, Marina Pasca Di Magliano

## Abstract

The adult healthy human pancreas has been poorly studied given lack of indication to obtain tissue from the pancreas in the absence of disease and rapid postmortem degradation. We obtained pancreata from brain dead donors thus avoiding any warm ischemia time. The 30 donors were diverse in age and race and had no known pancreas disease. Histopathological analysis of the samples revealed PanIN lesions in most individuals irrespective of age. Using a combination of multiplex immunohistochemistry, single cell RNA sequencing, and spatial transcriptomics, we provide the first ever characterization of the unique microenvironment of the adult human pancreas and of sporadic PanIN lesions. We compared healthy pancreata to pancreatic cancer and peritumoral tissue and observed distinct transcriptomic signatures in fibroblasts, and, to a lesser extent, macrophages. PanIN epithelial cells from healthy pancreata were remarkably transcriptionally similar to cancer cells, suggesting that neoplastic pathways are initiated early in tumorigenesis.

**Statement of significance:** The causes underlying the onset of pancreatic cancer remain largely unknown, hampering early detection and prevention strategies. Here, we show that PanIN are abundant in healthy individuals and present at a much higher rate than the incidence of pancreatic cancer, setting the stage for efforts to elucidate the microenvironmental and cell intrinsic factors that restrain, or, conversely, promote, malignant progression.

## Introduction

The pancreas, a main source of digestive enzymes, is uniquely susceptible to degradation *post mortem*(1-4). As there is no indication to sample the human pancreas in absence of a pathology, our understanding of this organ is limited, and has so far not benefited from advances in technology such as the single cell revolution of the last decade. Pancreatic ductal adenocarcinoma, the most common and deadly form of pancreatic cancer, arises from the exocrine pancreas(5,6). Different precursor lesions precede overt malignancy including mucinous lesions which can be seen on imaging and surgically treated as indicated(7-10). In contrast, pancreatic intraepithelial neoplasia (PanIN) is microscopic and typically only encountered following resection of other pancreatic pathologies, including overt cancer(11-13). Autopsy studies -mostly in an older patient population-indicate that PanIN is common in older individuals, however poor preservation of the pancreas due to prolonged warm ischemic time severely limits molecular analysis of autopsy samples(14-18). The prevalence of PanIN in the general healthy population remains unknown.

Pancreatic cancer is characterized by the accumulation of an extensive fibroinflammatory microenvironment, rich in extracellular matrix and including cellular components such as fibroblasts and immune cells. Genomic studies have characterized the mutation profiles of primary and metastatic tumors and reinforced the notion that the KRAS gene is almost invariably mutated in pancreatic cancer, together with common loss of tumor suppressor genes(19-22). Over the past several years, the advent of single cell technologies has given us unprecedented insight into the cellular composition of the tumor microenvironment, the heterogeneity across and within populations, and the signaling pathways driving cellular crosstalk(23-26). Pancreatic cancer is also characterized by accumulation of immune cells, including myeloid and lymphoid components; myeloid cells are largely immunosuppressive while T cells are exhausted in the majority of cancer patients(27-30). Whether corresponding populations of fibroblasts, myeloid cells, and T cells are present in the healthy human pancreas is unknown. A key limitation is that control samples are largely constituted by adjacent normal pancreas; while lacking malignant tumor cells, these samples are often not histologically normal and present with extensive inflammation and other alterations. PanIN studies, while revealing that these lesions share a similar mutation profile as cancer samples(10,31-36), are largely conducted in tissue from cancer patients or patients with different pancreatic pathologies, based on paraffin-stored samples. Thus, our understanding of the microenvironment of the human pancreas and of early lesions is extremely limited. While mouse models have been instrumental to understand the early stages of pancreas carcinogenesis and to study the healthy organ(37-40), they are a poor substitute for human samples. Mice are short-lived; laboratory mice spend their life in a highly controlled environment with specified diet, activity levels, light/dark cycle and none of the environmental and lifestyle stressors that characterize human life.

Here, we describe a unique partnership between the Pancreatic Disease Initiative at the University of Michigan and Gift of Life Michigan, a centralized organ procurement and allocation center located less than 6 miles from the University Hospital and its laboratories. While efforts are made to offer every organ to potential recipients in need, confounding factors often exist, including underlying disease of the donor (ie. diabetes and obesity) or, commonly, exhaustion of the recipient list which preclude transplantation, rendering organs available for research. This is particularly true for organs procured in donation after brain death (DBD). Unlike traditional organ donation where death occurs following cardiovascular decline and subsequent hypotension and hypoxia, DBD maintains blood flow to organs until they can be rapidly flushed with physiologic solution and immediately cooled to just above freezing. We posit these organs to be ideal for research analysis where limitation of hypoxia, electrolyte imbalance and rapid cooling will maximally preserve the cellular and transcriptomic profile of the normal pancreas. Our partnership with Gift of Life Michigan and close physical proximity has allowed for analysis of pancreata from 30 DBD donors of varied age, race and sex, resulting in a first ever comprehensive map of the microenvironment of the organ in humans. Additionally, because donors are brought to the Gift of Life Donor Care Center from across the state of Michigan, there is a broad range of ethnicity, socioeconomic status and environmental exposure magnifying the applicability of these findings.

## Results

### The adult human pancreas harbors frequent PanIN lesions

Through a collaboration with the Gift of Life Michigan Organ and Tissue program, we collected 30 pancreata from healthy adult organ donors, for whom no suitable recipient was identified. Of note, all donors were pre-screened via cross-sectional imaging and lab values (amylase and lipase) prior to organ recovery and were found to have no identifiable pathology in the pancreas. The organs were dissected from surrounding tissue and superior mesenteric and celiac arteries isolated. The arteries were cross-clamped and a cooled, isotonic physiologic solution was infused for 15 minutes. The organ was then rapidly removed and placed in sterile solution at 4°C and transported to the University of Michigan for processing within 90 minutes of cross clamp time (**Fig. 1A**). This method allows for no warm ischemic time, preserving the transcriptomic profile of the pancreas. As a pilot, we compared cell viability in samples from donation following brain death (DBD) with donation after circulatory death (DCD). In the latter, the heart and lungs are allowed to stop typically after a period of hypoxia and hypotension. This method is associated with a warm ischemic time of approximately 30-90 minutes resulting in a dramatic decrease in pancreatic cell viability of less than 20%; we thus discontinued the DCD program (**Supplementary FigS1A**). Patient age spanned from the third to eighth decade of life and included 20 males and 10 females. Our patient cohort included about 2/3 white donors, 1/3 African American donors, one donor of unknown race and, to date, one Asian donor, closely reflecting the demographic makeup of the State of Michigan (**Fig. 1B**, clinical table in **Supplementary Table 1**). For each sample, we collected 15 tissue blocks from multiple regions along the head-tail axis **(Supplementary FigS1B)**. Histopathological analysis of each block revealed PanINs in 18 of 30 donor specimens (**Fig 1B** and **1C**) as well as normal acinar parenchyma, normal ducts and areas of acinar-to-ductal metaplasia (ADM) (**Supplementary FigS1C**). PanIN distribution ranged from isolated lesions in one region of the pancreas to multifocal. PanINs were surrounded by an area of fibrosis with interspersed cellular components (**Fig 1C**).

**Fig 1.**
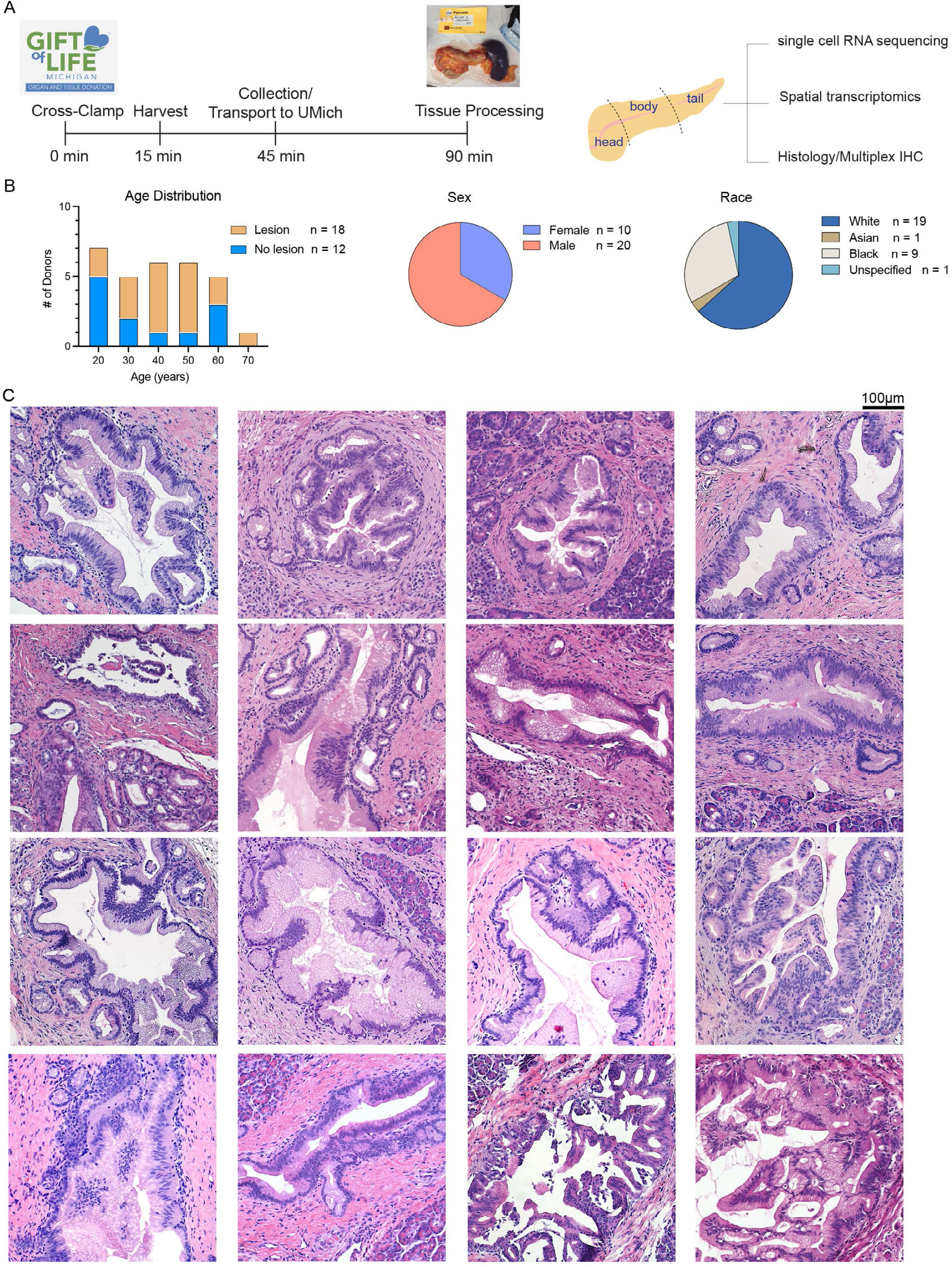
The adult human pancreas harbors frequent PanIN lesions. A) Schematic of workflow to recover and process Gift of Life donor pancreas organs. B) (Left) Population distribution barplot of donor cohort. Tan bars represent donors that were found to have PanIN lesions on histological examination. Blue bars represent donors in whom no neoplastic lesions were found. (Middle) Pie chart of donor cohort by sex. (Right) Pie chart of donor cohort by race. C) H&E of representative PanIN lesions found in donor pancreata. Each H&E section represents a different donor.

### PanINs from healthy individuals are surrounded by a unique microenvironment

To characterize the immune and stromal composition of the PanIN microenvironment, we performed multiplex fluorescent immunohistochemistry on formalin-fixed, paraffin-embedded tissue. We used a previously described immune panel (24) (**Fig 2A, Supplementary FigS2A**) and a newly optimized fibroblast panel (**Fig 2B, Supplementary FigS2B**). We then quantified cellular composition in areas including PanIN compared with areas including normal acini, ducts or ADM. Myeloid cells were enriched in ADM and PanIN areas compared to areas surrounding normal acini and ducts; T cells were almost exclusively detected surrounding PanINs when compared to any other histological area (**Fig 2C**). Abundant fibroblasts surrounded both normal ducts and PanINs, although the latter showed more heterogeneous staining for vimentin, αSMA, PDGFRb, and FAP (**Fig 2B**), possibly indicating higher complexity of subtypes. Thus, PanIN formation is accompanied by the establishment of a unique microenvironment rich in fibroblasts, myeloid and T cells, distinct from the histologically normal pancreas.

**Fig 2.**
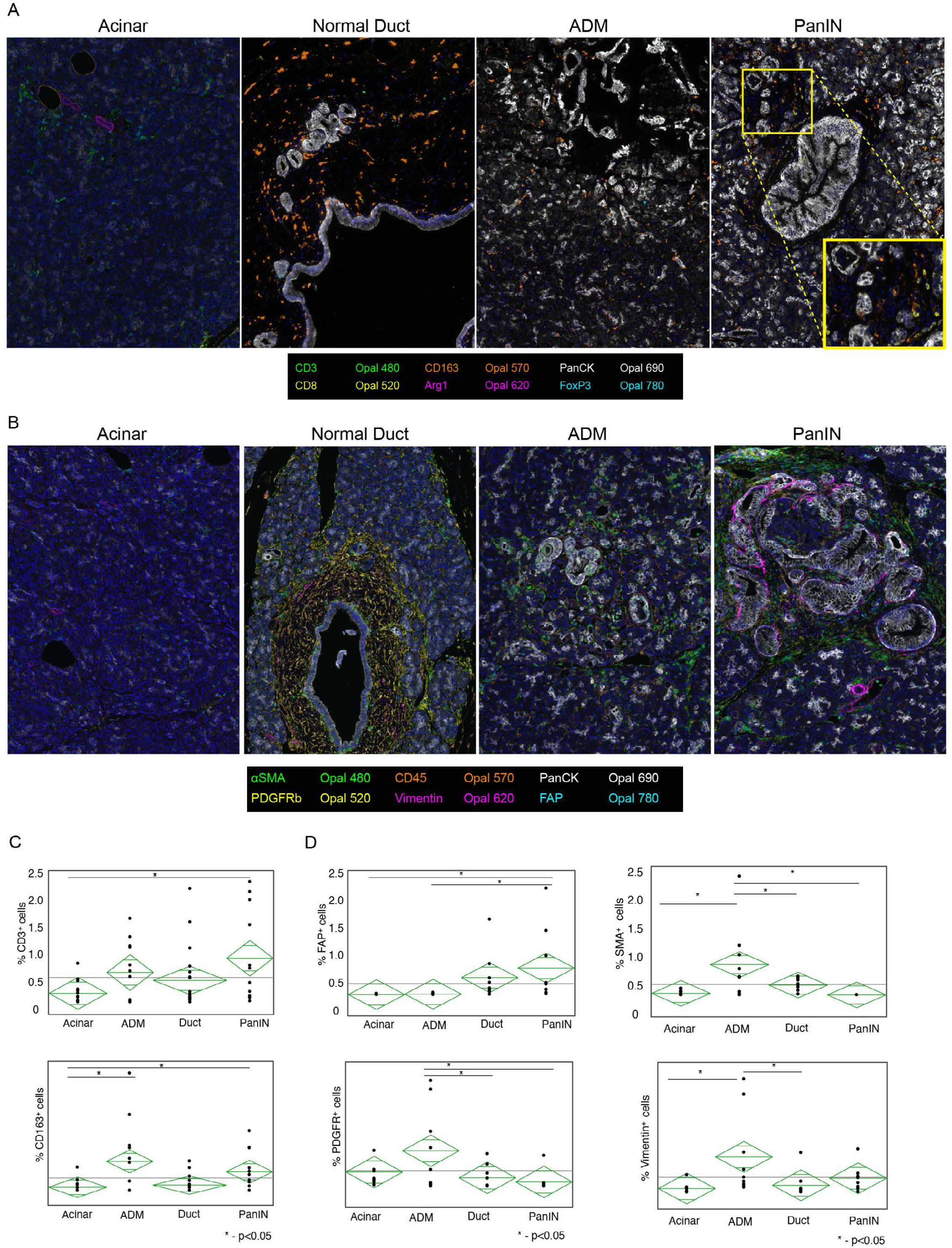
PanIN lesions in healthy pancreata is surrounded by a unique microenvironment. A) mfIHC composite images of formalin-fixed, paraffin-embedded donor tissue specimens, highlighting acinar, normal duct, ADM (acinar-to-ductal), and PanIN structures. Antibodies and colors of immune panel used in the legend below. B) mfIHC composite images of formalin-fixed, paraffin-embedded donor tissue specimens, highlighting acinar, normal duct, ADM (acinar-to-ductal), and PanIN structures. Antibodies and colors of fibroblast panel used in the legend below. C) Quantification of percent positive CD3+ cells (top) and CD163+ cells (bottom) surrounding Acinar, ADM, Duct, and PanIN populations, respectively. Asterisks denote a P value of <0.05, as determined by ANOVA. D) Quantification of percent positive FAP^+^ cells (top left) and FDGFR^+^ cells (bottom left), SMA^+^ cells (top right), and Vimentin^+^ cells (bottom right) surrounding Acinar, ADM, Duct, and PanIN populations, respectively. Asterisks denote a P value of <0.05, as determined by ANOVA.

To gather further insight in the composition of PanIN and its microenvironment, we performed single cell RNA sequencing of 6 donor organs, for a total of 44,068 cells (**Fig 3A**). For five of the pancreata, we sequenced head and tail samples separately **(Supplementary Table 1**, last column). On histological analysis, 4 of the pancreata had PanIN lesions while no lesions were detected in the remaining two. We used uniform manifold approximation and projection (UMAP) for visualization of cell populations and identified each population based on published lineage markers, detecting epithelial cells (acini and ducts) as well as non-epithelial cells (myeloid cells, lymphocytes, fibroblasts, and endothelial cells) (**Fig 3A, Supplementary FigS3A**). The cellular composition was variable across samples, with no significant trend based on whether individual pancreata harbored lesions (**Fig 3B**).

**Fig 3.**
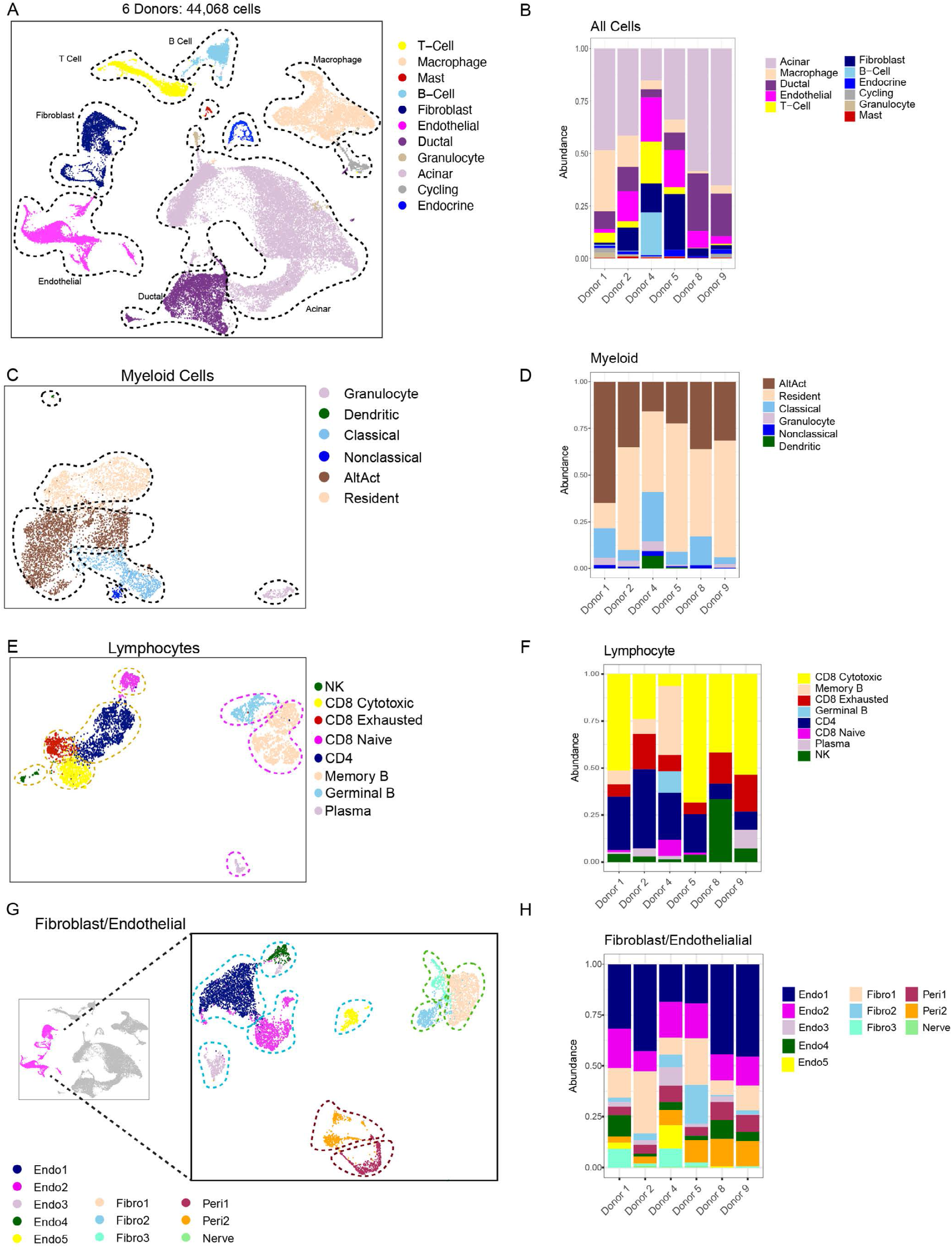
Healthy pancreata contain several non-epithelial populations, including myeloid, lymphocyte, fibroblast, and endothelial cells. A) UMAP of all cells captured from single cell RNA sequencing of six donor pancreata. Populations are identified by color. B) Histogram of cell type abundance of all captured cells by donor. C) UMAP of extracted myeloid cells from donor pancreata. Populations are identified by color. AltAct = Alternatively Activated Macrophages. D) Histogram of cell type abundance of specific myeloid cell populations by donor. E) UMAP of extracted lymphocytes from donor pancreata. Populations are identified by color. F) Histogram of cell type abundance of specific lymphocyte populations by donor. G) UMAP of extracted fibroblast, pericyte, and endothelial populations from donor pancreata. Populations are identified by color. H) Histogram of cell type abundance of specific fibroblast, pericyte, and endothelial populations by donor.

To evaluate the spectrum of myeloid populations in the human pancreas, we subsetted the myeloid cells from the single cell dataset (**Fig 3C**) and identified subclusters based on previously published signatures (**Supplemental FigS3B**) (23). Myeloid populations were largely dominated by alternatively activated, resident and classical macrophages, while granulocytes were rare to undetectable (**Fig 3C, Fig 3D)**. Alternatively activated macrophages expressed the expected markers of *MARCO* (scavenging receptor), *APOE*, and *SPP1*; resident macrophages were high in complement component encoding genes; classical macrophages expressed *CCR2, LYZ*, and, *FCN1;* finally, *a* very small population of nonclassical (*CD14*^*Low*^) macrophages expressed *FCGR3A* and *LILRA5* (**Supplementary FigS3C**).

Pancreata also harbored lymphocytes, including CD4^+^ and CD8^+^T cells, B cells, and NK cells (**Fig 3E, Fig 3F**). We did not readily capture any regulatory T cells. We detected both cytotoxic and exhausted CD8^+^ T cells, the former identified by *KLRC* and *XCL1* expression; the latter by *GZMK* and *KLRG1* expression (**Supplementary FigS4A, FigS4B**). Analysis of B lymphocytes showed a large population of memory B cells expressing *IGHM* and *IGHD*, germinal B cells marked by *RGS13* expression, and plasma cells characterized by lack of *HLA-DRA* and presence of *IGHG4, SDC1, and IGHG1* (**Supplementary FigS4A, FigS4C**).

Thus, the human pancreas houses a variety of innate and adaptive immune cells. Of note, the dissociation step, required for single cell RNA sequencing, precludes determination of whether any of the immune cells were derived from normal pancreas parenchyma, or from areas surrounding PanIN lesions; however our multiplex immunostaining data shows that macrophages are present throughout the tissue while T cells are more likely to localize near PanIN (**Fig 2)**.

We then analyzed fibroblasts, endothelial cells, and pericytes together, as these cells share several markers. We identified 3 distinct populations of fibroblasts, Fibro1 (*DPT* low), Fibro 2 (*DPT* high), and Fibro3 (expressing *FBLN2* and *CFD*) (**Fig 3G-H, Supplemental FigS5A**). Notably, *DPT* was recently described as a pan-fibroblast marker across multiple organs and diseases states in mouse tissues(41); based on our data, *DPT* is expressed across fibroblast populations in the adult human pancreas, albeit at variable levels (**Supplemental FigS5B)**. Fibroblasts also had universally high expression of *PDGFRB/A*, and, to a lesser extent, *PDPN* (**Supplemental FigS5B-C**). *PDGFRB* was also expressed by pericytes while *PDGFRA* was only expressed in fibroblasts. In contrast, *ACTA2* was expressed by a small subset of fibroblasts only (**Supplemental FigS5B**). *LRRC15*, a marker recently associated with tumor promoting pancreatic CAFs (42), was not expressed in normal pancreas fibroblasts, but was detected in a few cells in the Endo5 endothelial cell cluster(**Supplemental FigS5B**). Lastly, FABP4, which labels pancreatic stellate cells in mice (43), only labeled a few fibroblasts but was widely expressed by pericytes and endothelial cells, possibly highlighting a mouse/human difference or reflecting the age of the donors (**Supplemental FigS5B**). We identified 5 endothelial cell populations (Endo1-5, **Figure 3G-H, Supplemental Figure S5A and S5D**), expressing varying levels of *VWF, CDH5, FABP5*, and *CD74* (**Supplemental FigS5B, FigS5D**), and two pericyte populations, expressing RGS5 (**Supplemental FigS5A**). The Endo5 population was notable given its elevated expression of angiogenic factors *IL33, TFF3, CXCL2*, and *KRT18* (**Supplemental FigS5A, Supplemental FigS5D**). The prevalence of each cell type, both for immune and non-immune components of the microenvironment was variable across donor samples.

Overall, the normal pancreas includes a heterogeneous set of structural cells, such as fibroblasts, pericytes, and endothelial cells, as well as immune cells. The normal human pancreas also presents with histological heterogeneity reflecting age and the complexities of human lifestyle and genetics.

### Comparison of the microenvironment in healthy pancreata and pancreatic tumors reveals distinct stromal features

We next sought to investigate the differences in the microenvironment of the normal pancreas and pancreatic cancer. For this purpose, we integrated single cell RNA sequencing data from healthy donor samples (6 pancreata; 11 samples total) with our previously published dataset(44) of tumor (n=16) and adjacent normal (n=3) samples. The latter were obtained from patients undergoing surgery for duodenal adenoma, ampullary carcinoma or PDA, respectively, and tissue was confirmed to be free of cancer by pathologic evaluation. UMAP two-dimensional visualization showed a wide variety of epithelial and non-epithelial cells (**Fig 4A**). In terms of cell type composition, each sample was unique (**Supplemental FigS6A**); however, some common themes emerged. As expected, acinar cells were captured in abundance in the normal samples and, to an extent, in adjacent normal, while they were rarely captured in tumor samples (**Fig 4B** and **Supplemental FigS6B**). Conversely, tumor samples had a higher proportion of macrophages and CD4^+^ T cells. Granulocytes were rarely detected in normal samples and only observed in 1 out of 3 adjacent normal samples, but they were abundant in most tumor samples. The proportion of CD8^+^ T cells was variable across each group of samples. Endothelial cells were generally more abundant in healthy pancreata and adjacent normal, although occasionally detected in individual tumors. Using differential abundance analysis to generate a neighborhood graph differential abundance plot (45) between healthy and tumor samples, we defined cell populations with significant shifts in abundance between tumor and normal states (**Fig 4C)**. We generated a beeswarm differential abundance plot(45) and found that across all samples, comparing tumors vs. normal showed loss of acinar cells and endothelial cells, and an increase in myeloid cells and non-acinar epithelial cells, likely cancer cells, in the tumor samples (**Fig 4D)**.

**Fig 4.**
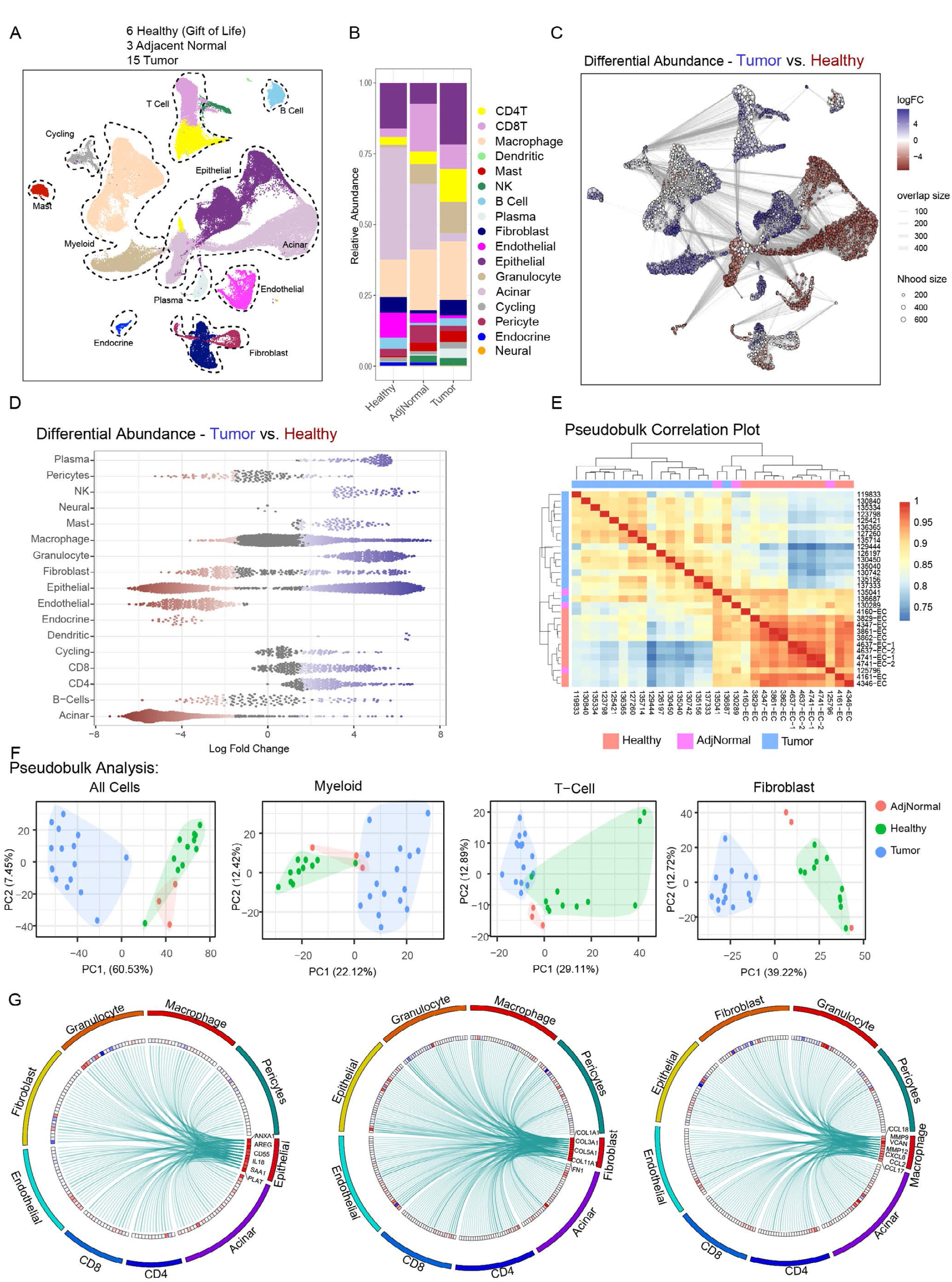
Comparison of the microenvironment in healthy pancreata and pancreatic tumors reveals distinct stromal features. A) UMAP of all cells captured from single cell RNA sequencing of six donor pancreata merged with 15 PDAC samples and 3 adjacent normal samples. Populations are identified by color. B) Histogram of cell type abundance of all cell populations by disease state (healthy, adjacent normal, and tumor). C) Neighborhood graph differential abundance plot of merged tumor, healthy, and adjacent normal samples. Size of dots represent neighborhoods, while edges represent depict the number of cells shared between neighborhoods. Neighborhoods colored in red represent significantly increased abundance in healthy samples while neighborhoods colored in blue represent significantly increased abudance in tumor samples. D) Beeswarm plot of differential abundance by cell type. X-axis represents log-fold change in abundance between tumor and healthy states. Each dot is a neighborhood; neighborhoods colored in red represent significnatly increased abundance in healthy samples while neighborhoods colored in blue represent signficantly increased abudance in tumor samples. E) Correlation heatmap of pseudobulk-aggregrated counts of 15 tumor samples, 3 adjacent normal samples and 11 donor samples (given that single cell sequencing was performed on the head and tail sections separately of 5 out of 6 donors). Each row/line represents one aggregated single cell sequencing sample. F) PCA plots of pseudobulk-aggregated counts from all cells, myeloid cells, t-cells, and fibroblasts. Each dot represents one aggregated single cell sequencing sample. G) Circos plots of putative ligand–receptor interactions that are upregulated in PDAC epithelial (left), fibroblast (middle), and macrophage (right) cells compared with healthy cells. The heatmap within the circos plots is the scaled average expression of each gene within PDAC tissue cell populations. The interactions plotted are those in which the expression level of the ligand is increased in PDAC samples compared with healthy tissues.

To determine whether gene expression alone could distinguish between tumor samples and normal pancreas, we aggregated sequencing data from all of the cells in each sample (pseudo-bulk) and applied correlation and principal component analysis (PCA) on normalized gene-sample count matrices. Both plots showed that tumor samples largely clustered separately from healthy samples; adjacent normal samples were variable (**Fig 4E, 4F**). The classification of adjacent normal is to be interpreted with caution given the relatively small sample size. As pseudobulk analysis of total cells is affected by cellular composition in each sample, we then repeated analysis on a population-specific basis, initially focusing on broad stromal cell populations, namely myeloid cells, T cells and fibroblasts. The PCA plots revealed clear clustering of tumor vs normal samples, while adjacent normal were variable, and at times interspersed with the healthy and/or tumor samples (**Fig. 4F**). Lastly, we plotted predicted Ligand-Receptor interactions(46) between cell populations that were increased in tumors compared with normal samples (**Supplementary Table 2)**. Epithelial signals enriched in tumor samples included *EGFR* ligands, consistent with activation of this pathway in pancreatic cancer (47,48). *SAA1*, a factor linked to promotion of pancreatic carcinogenesis (49), was also elevated in tumors (**Fig 4G**). Signaling from fibroblasts (as well as from pericytes and endothelial cells) was prominently increased in tumor samples when all possible interactions were measured (**Supplemental Figure S6C, S6D**); specific findings includes an increase in extracellular matrix signaling driven by collagen and fibronectin (**Fig 4G**), consistent with the fibrotic microenvironment of pancreatic cancer (50). Macrophages showed an increased production of *CCL* and *CXCL* family cytokines, linked to tumor-promoting and immunosuppressive pathways (51,52).

To further investigate changes in cellular composition and in gene expression patterns, we further analyzed specific cell populations For this comparison, we focused on myeloid cells and fibroblasts, cell types that were present in abundance both in tumor and normal samples. Myeloid cells included granulocytes, derived largely from tumors, and several populations of macrophages detected across samples (**Fig 5A, Supplemental Fig 7A**). Dendritic cells (including FCER1A^+^, CLEC9A^+^, and LAMP3^+^ DCs) were detected in a few of the samples, but not abundant enough for meaningful gene expression comparisons. In contrast alternatively activated, classical, nonclassical and resident macrophages (defined based on the markers shown in **Supplemental FigS7B**) were abundant both in the healthy pancreas and in pancreatic cancer. PCA analysis of each of these populations revealed distinct gene expression patterns in tumors vs healthy organs; adjacent normal were again variable and interspersed with healthy and tumor samples (**Fig 5B**). Pathway annotation revealed an increase in chemotaxis pathways in tumors vs healthy pancreata, indicating that signaling programs in myeloid migration as largely responsible for the transcriptomic differences between tumor and healthy states (**Supplemental FigS7C**). Performing differential expression analysis between macrophages from the healthy organs versus tumors, interestingly, did not reveal any signature markers of tumor associated macrophages (53) (**Fig 5C**). We then individually plotted known TAM markers. Among those, *APOE, MRC1, C1QA, C1QB* and *SPP1*, described by our group and others as TAM-specific in mouse models (53-55), were similarly expressed in healthy pancreas-associated macrophages as well as TAMs (**Fig 5D)**. This finding was surprising and differentiated our human data from mouse model findings, where all of these factors are lowly expressed or undetectable in healthy organs.

**Fig 5.**
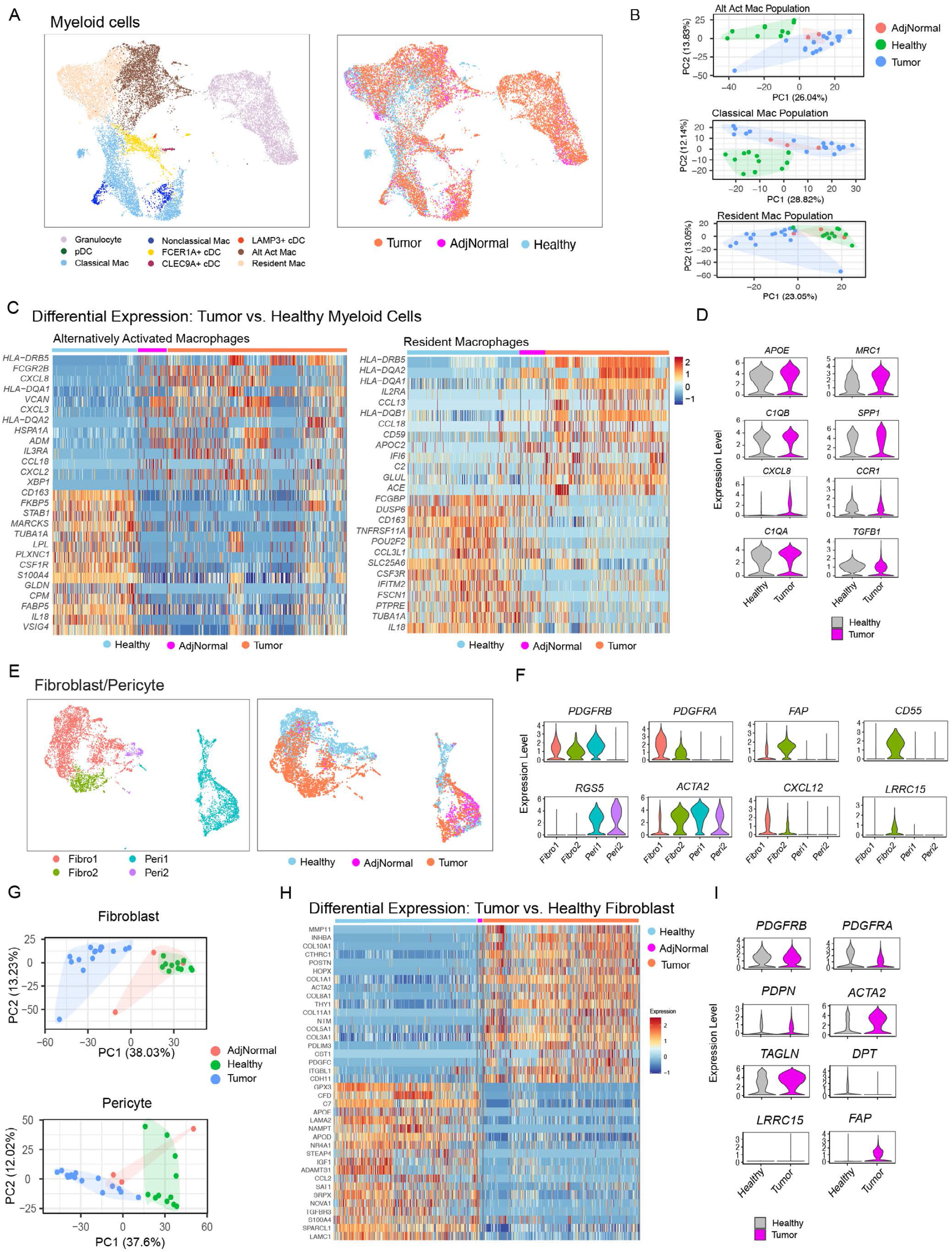
Myeloid and fibroblast populations from tumor-bearing pancreata display distinct transcriptomic signatures compared to their non-tumor counterparts. A) (left) UMAP of extracted myeloid cells from single cell dataset of healthy, adjacent normal, and tumor samples. Populations are identified by color. AltAct = Alternatively Activated Macrophages. pDC = Plasmacytoid Dendritic Cells. cDC = Conventional Dendritic Cells. (right) UMAP overlay of disease states on extracted myeloid cells from single cell dataset of healthy, adjacent normal, and tumor samples. B) PCA plots of pseudobulk-aggregated counts from specific myeloid cell populations Each dot represents one aggregated single cell sequencing sample. C) Top differentially expressed genes between alternatively activated macrophages (left) and resident macrophages (right) from healthy (blue) and tumor (orange) samples. D) Violin plots of normalized expression of select tumor-associated macrophage markers comparing healthy to tumor samples. E) (left) UMAP of extracted fibroblast and pericyte cells from single cell dataset of healthy, adjacent normal, and tumor samples. Populations are identified by color. (right) UMAP overlay of disease states on extracted fibroblast/pericyte cells from single cell dataset of healthy, adjacent normal, and tumor samples. F) Violin plots of normalized expression of select fibroblast markers mapped across fibroblast and pericyte populations. G) PCA plots of pseudobulk-aggregated counts from fibroblast (top) and pericyte (bottom) populations Each dot represents one aggregated single cell sequencing sample. H) Top differentially expressed genes between fibroblasts from healthy (blue) and tumor (orange) samples. D) Violin plots of normalized expression of select fibroblast markers comparing healthy to tumor samples.

Fibroblast and pericyte clustering identified two distinct fibroblast and two distinct pericyte populations (**Fig 5E-F** and **Supplemental FigS7D-E**). Fibroblast cluster Fibro2 was almost entirely derived from tumor samples, while fibroblast cluster Fibro1 was derived from both healthy and tumor samples, predominantly the former. Interestingly, fibroblast cluster Fibro2 expressed relatively high levels of the myCAF(23) marker *ACTA2* and decreased expression of the iCAF(23) marker *CXCL12*, in addition to increased expression of FAP (a cancer-associated fibroblast marker known to mediate immunosuppression(56)) and exclusive expression of CD55 and LRRC15. These data show that healthy pancreas fibroblasts are altogether distinct from cancer-associated fibroblasts, a notion supported by distinct clustering on PCA analysis(**Fig 5G**). Differential expression analysis again revealed an increase in activation markers and markers of tumor promoting fibroblasts in tumor samples, such as ACTA2, FAP and LRRC15 (**Fig 5H** and **5I**).

Of note, we also captured several lymphocyte and NK populations, although this group derived from tumor samples, with the exception of naïve T cells, which were mostly detected in healthy organs (**Supplemental FigS8A** and **S8B**). PCA plots revealed a spectrum, with some tumor T cells clustering within the normal T cell populations, and others quite distinct (**Supplemental FigS8C**); pathway analysis showed tumor samples were enriched for genes in negative regulation of T cell proliferation (**Supplemental Fig 8D**). However, overall these T cells were less distinct between tumor and normal, an interesting finding given the hallmarks of poor immunogenicity, immune exhaustion, and impaired cytotoxic activity of T cells in pancreatic cancer (24,57).

Overall, our analysis of human pancreata shows frequent PanINs in organs from healthy individuals. Comparison with PanINs with tumors samples identified shared inflammatory gene expression profiles of macrophages. In contrast, fibroblasts were clearly divergent in tumors compared to PanIN. Finally, T cells were rare to undetectable in normal areas of the pancreas, present in small numbers in PanINs and common in pancreatic cancer, although subsets such as Tregs were almost uniquely present only in tumors. Thus, comparative analysis reveals both similarities and differences in the components of the microenvironment.

### Spatial transcriptomics reveals a unique epithelial gene signature of PanIN lesions which aligns closely with tumor epithelium

We next focused on the epithelial population in our single cell RNA sequnecing dataset to determine if we could identify transitory cell populations, including cells undergoing acinar to ductal metaplasia and cells from PanIN lesions. Analysis of the epithelial and acinar populations revealed 16 distinct subclusters (**Supplemental FigS9A, S9B**), with clusters 4, 5, and 6 derived mainly from tumor samples(**Supplemental FigS9C**). These clusters had high expression of *S100* genes, encoding for a family of proteins often elevated in cancer cells (58), and linked to pancreatic cancer metastasis (59) (**Supplemental Fig 9D**). In the quest to identify any PanIN cells in donor samples, we investigated previously reported transcriptomic signatures specific to acinar, ductal, PanIN, and cancer cells (60) in our sequencing data. Using enrichment analysis, we defined clusters enriched for the acinar, ductal, and cancer cell signatures. As expected, the acinar signature marked cells largely derived from healthy and adjacent normal samples, while tumor signature marked cells from PDAC samples (**Supplemental FigS9E**). We also identified cell populations enriched for the previously described Duct-like1 (expressing normal ductal genes) and Duct-like2 (with increased expression of mucus and trefoil factor genes) signatures. In our dataset, the former mapped to cells from normal and adjacent normal samples, the latter mapping within tumor-derived cells **(Supplemental FigS9E)**. In contrast, we were unable to map the PanIN signature in our single cell dataset (**Supplemental FigS9E**). Interestingly, utilizing genes for *HALLMARK_KRAS_SIGNALING_UP* and *_DOWN(61)*, we observed tumor-derived epithelial cells were relatively enriched for genes upregulated with KRAS activation and relatively depleted from genes downregulated with KRAS activation (**Supplemental FigS9F**). We hypothesized that PanIN cells, a relatively rare population, were not captured in our single cell dataset; another possibility is that the PanIN signature derived from tumor adjacent PanIN, as in previous studies (60), is distinct from sporadic PanIN in healthy pancreata.

To directly evaluate the gene expression pattern of PanIN, we utilized the GeoMx Nanostring spatial transcriptomics platform. After careful review of tissue slides with a clinical pathologist (JS, who specializes in pancreatic histopathology), acinar, normal duct, ADM, and PanIN regions of interest (ROIs) were selected from the donor pancreata, while tumor-associated PanINs, glandular tumors, and poorly differentiated tumor ROIs were selected from malignant samples (**Fig 6A-B**). In addition, we collected ROIs from acini and normal ducts adjacent to tumors for comparison. To avoid confounding transcripts from the stroma, samples from each region were segmented using pancytokeratin (PanCK) and CD45 to select PanCK^+^/CD45^-^ cells in all ROIs, with the exception of acinar ROIs, where we selected PanCK^-^/CD45^-^ (as acinar cells do not express PanCK, and can be easily identified histologically). Following quality control and data normalization, we performed PCA analysis, and detected a prominent batch effect. Upon correction, we obtained acinar and PanIN ROIs that clustered separately, while ADM and normal duct ROIs were interspersed with each other, indicating transcriptional similarity, which is expected (**Supplemental FigS10A**). To extract gene signatures specific to each cell type, we applied linear mixed model differential expression analysis comparing each cell type to all other remaining cell types (**Supplemental FigS10B, Fig 6C, Supplemental Table 3**). We thus identified expected digestive enzymes in acinar ROIs (*CEL, CPA1, CELA2B, PRSS3*), and genes encoding for mucins in ductal ROIs (*MUC5B, MUC20*). Ductal ROIs also expressed *AQP3* ⎯encoding for a water channel protein typically found on the basolateral membrane of ducts(62) ⎯, *MUC5B* ⎯ expressed in noncancerous pancreatic ductal epithelium(63) ⎯ and *MUC20*, which expression has not been previously reported in the pancreas. PanIN ROIs were enriched for the previously described marker *CLDN18(60,64)*. We also detected expression of trefoil factor genes *TFF1* and *TFF2*, secretory proteins expressed in PanINs and, to a lesser degree, in early PDAC(65,66). Interestingly, *MUC5AC*, a known PanIN marker(67), was not specifically enriched in PanIN, as its expression was higher in glandular tumors and ducts (**Supplemental FigS10C)**. ADM ROIs included *AQP1* (in contrast to normal ducts marked by *AQP3*), as well as serpin family genes (*SERPING1)* and complement (*C6*), an interesting combination as serpins regulate the complement cascade(68). While the complement system has been implicated in the pathogenesis of pancreatic cancer(69,70), this is the first report of its upregulation in ADM from healthy pancreas, suggesting that complement dysregulation may be an early pre-neoplastic event. Lastly, glandular and poorly differentiated tumor ROIs were distinct, with glandular tumors more similar to PanIN, as expected. While healthy samples and tumor-bearing samples had a high degree of concordance in marker features for acinar, ductal, and PanIN marker features (as ADM was only captured in healthy samples), we noted that PanINs from tumor-bearing samples had higher expression of genes seen common in tumor ROIs, suggesting closer association with malignancy.

**Fig 6.**
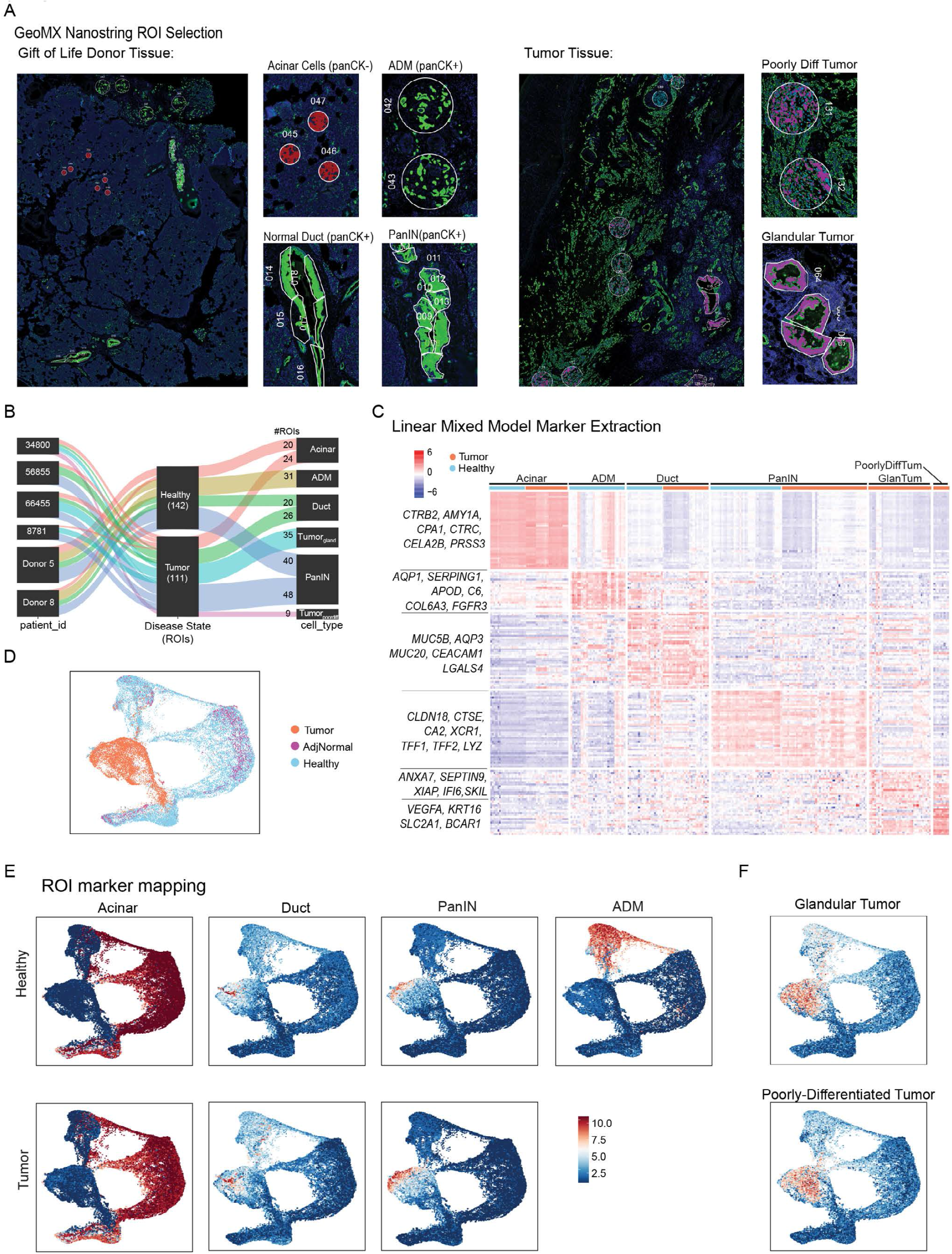
Spatial transcriptomics reveals a unique epithelial gene signature of PanIN lesions which aligns closely with tumor epithelium. A) (Left) GeoMX tissue section from donor pancreas, stained for panCK and CD45. panCK^+^ segments were pseudo-colored in green, while panCK^-^CD45^-^ segments were pseudo-colored in red. (Right) GeoMX tissue section from surgically resected treatment-naïve PDAC, stained for panCK and CD45. panCK^+^ segments were pseudo-colored in purple. B) Sankey plot showing distribution of ROIs amount donor or PDAC samples. C) Heatmap of cell-type specific markers derived from differential gene expression using the linear mixed model on spatial transcriptomic ROIs. D) UMAP overlay of disease states on extracted epithelial cells from single cell dataset of healthy, adjacent normal, and tumor samples. E) AUCell geneset scoring mapped to epithelial single cell dataset of healthy, adjacent normal, and tumor samples using signatures derived from Acinar, Normal Duct, PanIN, and ADM (acinar to ductal metaplasia) spatial transcriptomic ROIs. Top row represents spatial transcriptomic signatures obtained from healthy tissue; bottom row represents spatial transcriptomic signatures obtained from tumor tissue. F) AUCell geneset scoring mapped to epithelial single cell dataset of healthy, adjacent normal, and tumor samples using signatures derived from glandular tumor and poorly differentiated tumor ROIs.

Using Gene Set Variation Analysis (GSVA), we investigated the enrichment of the previously reported PDAC subtype-specific signatures (71-73). While acinar and ADM ROIs showed relative enrichment of ADEX and exocrine-like subtypes signatures, PanIN ROIs were enriched to classical signature, and the poorly differentiated tumor ROIs displayed basal signature enrichment. Interestingly, PanIN ROIs corresponded more closely to the classical signature than glandular tumors, as the latter showed expression of basal-like genes as well, consistent with recent reports on intermediary states between classical and basal in most human tumors at the single cell level(26,74) (**Supplemental Fig. S10D**). In concordance with our prior differential expression analysis showing differences between PanINs from different sources, we noted that PanINs from tumor-bearing samples expressed some basal-like genes whereas PanINs from healthy samples showed did not (**Fig 6C**). Thus, sporadic PanINs from otherwise healthy pancreata are similar, but distinct, from tumor adjacent PanINs.

We then integrated spatial transcriptomics with single cell RNA sequencing data. Using the top 20 genes from each ROI class, we defined cell-type specific gene signatures for acinar, ductal, ADM, PanIN and tumor cells and mapped these signatures to the epithelial cluster from the single cell epithelial sequencing data (**Fig 6D-F**). Our aim was to map cell populations relatively enriched for each defined signature. As expected, the spatial-transcriptomic derived signatures identified distinct acinar, ADM, and ductal populations (**Fig 6E**). The glandular and poorly differentiated tumor signatures were, as expected, enriched in cells derived from tumor samples (**Fig 6F**). While the glandular tumor signature mapped to cells spanning most of the epithelial population, consistent with its heterogeneous subtype scoring, the poorly differentiated tumor signature was confined to tumor-derived epithelial cells. Interestingly, the ADM signature was enriched in the donor-derived epithelial cells, while the PanIN and normal ductal signatures from both healthy and tumor-bearing samples only mapped to specific clusters derived from tumor samples, an unexpected finding.

PanINs from tumors notably marked a slightly larger population of cells within the tumor-derived epithelial population than PanINs from healthy samples. This supported the differential expression analysis, which showed more shared features of tumor-bearing PanINs ROIs with tumor ROIs, compared with PanINs from healthy samples.

Overall, sporadic PanINs from healthy organs were surprisingly similar to tumor epithelial cells at the transcriptional level, suggesting that these neoplastic lesions have already acquired many features of malignant cells.

### Claudin 18, MUC5AC/B, and AQP1/3 distinguish normal Ducts, ADM, and PanIN in healthy pancreas

We next endeavored to validate our gene signatures in donor tissue to determine whether they accurately identified different epithelial populations, as suggested by our transcriptomic data. We selected markers that distinguished each epithelial cell type (**Fig 6C**). Our list included AQP1 (ADM-specific), MUC5B and AQP3 (normal-duct-specific), and CLDN18 and and *in situ* hybridization probe for *TFF1*(PanIN-specific), for which antibodies were readily available. We included MUC5AC as a PanIN marker, as our data supports its exclusive expression in PanINs in non-tumor bearing samples (compared to normal ducts, ADM, and acinar cells) **(Supplemental FigS10C)**. For each of these markers, we performed co-immunofluorescent staining on sections obtained from 4 different donor pancreata, including E-cadherin as a common epithelial lineage marker (**Supplemental FigS11-14)**. We also stained for p-ERK to assess the status of ERK activation in normal pancreas within acinar cells, ADM, normal ducts, and PanINs. In sections containing both PanIN and normal duct, MUC5B only stained normal ducts, while MUC5AC was specific for PanIN (**Fig 7A** and **7B**). CLDN18 and pERK staining was elevated in PanIN (**Fig 7C** and **7D**). AQP3 only stained normal ducts, while AQP1 was specific to ADM (**Fig 7E** and **7F**). Lastly, *TFF1* was co-expressed with CLDN18 in PanIN (**Fig 7G-H)**. The expression pattern held thru across samples (**Supplemental FigS11-14**, each showing consecutive serial sections).

**Fig 7).**
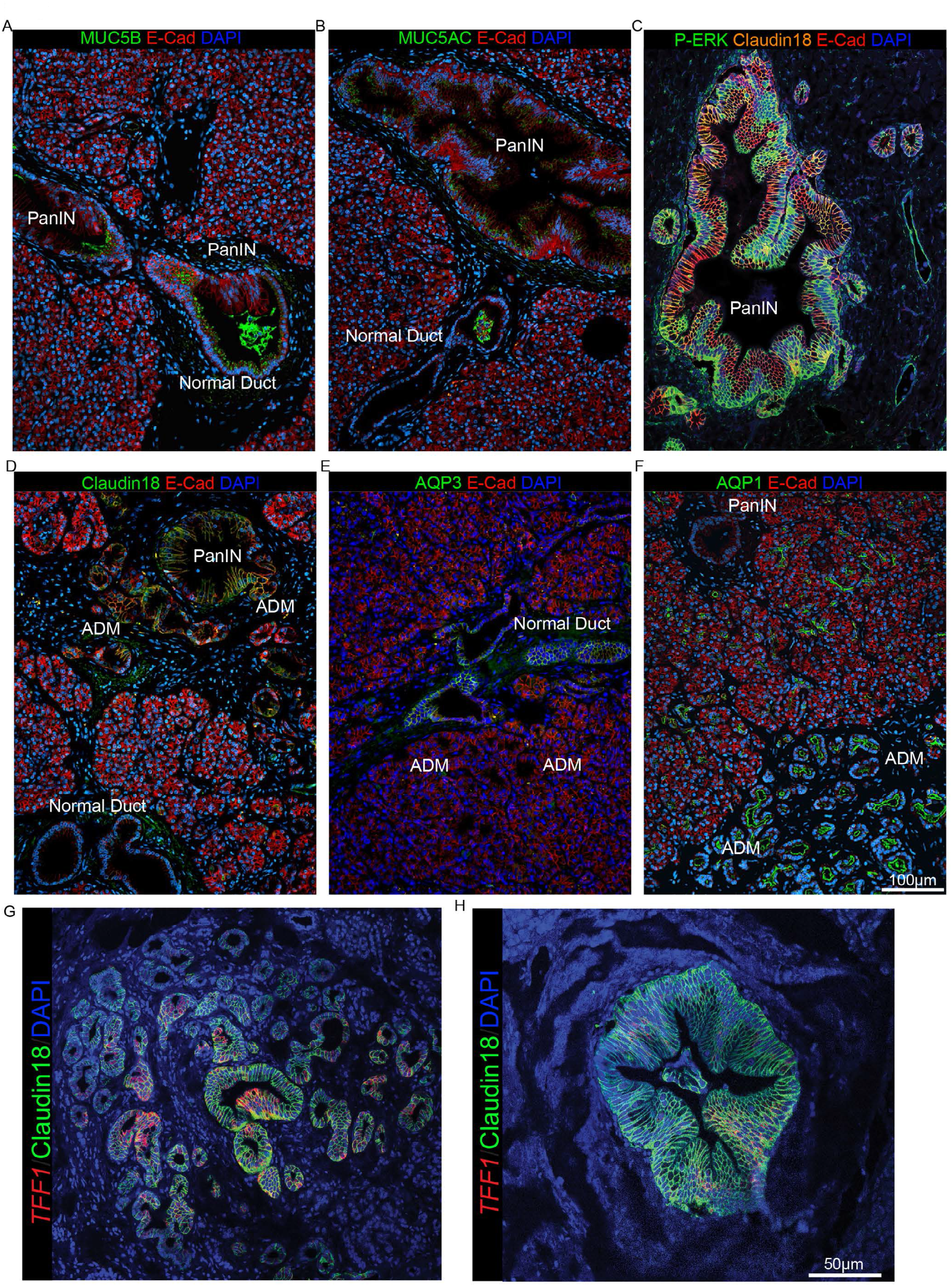
Claudin 18, MUC5AC/B, and AQP1/3 distinguish normal Ducts, ADM, and PanIN in healthy pancreas. A) Donor tissue stained with antibodies against Muc5B (green) and E-Cad (red), along with DAPI. B) Donor tissue stained with antibodies against Muc5AC (green) and E-Cad (red), along with DAPI. C) Donor tissue stained with antibodies against p-ERK (green), Claudin18 (orange), and E-Cad (red), along with DAPI. D) Donor tissue stained with antibodies against Claudin18 (green) and E-Cad (red), along with DAPI. E) Donor tissue stained with antibodies against Aqp3 (green) and E-Cad (red), along with DAPI. F) Donor tissue stained with antibodies against Aqp1 (green) and E-Cad (red), along with DAPI. G-H) Donor tissue stained with RNAScope probe for *TFF1* (red) and antibody against Claudin18 (green), along with DAPI.

Thus, spatial transcriptomics-derived signatures defined individual epithelial cell populations and correctly predicted protein expression of unique lineage markers.

## Discussion

The pancreas and its constitutive acinar cells possess high metabolic activity and undergo rapid cell death under hypoxic conditions(1-4). During this process, granules containing proteolytic and hydrolyzing enzymes are released resulting in rapid destruction of cellular components and nucleic acids. As such a true transcriptomic profile of the physiologically normal pancreas has remained elusive. Due to rapid degradation postmortem, autopsy samples are not suitable for techniques requiring intact RNA, including single cell RNA sequencing, and are often difficult to examine by immunostaining as well. The mouse pancreas, while histologically similar to the human counterpart, does not recapitulate the effects of age and environmental stressors that characterize human life.

As a result, single cell studies of the human pancreas have included very few samples(75,76) or embryonic samples(77) and have focused on endocrine cells less susceptible to degradation. Previous autopsy studies on pancreata of patients deceased with no known pancreas pathology revealed frequent PanINs and KRAS mutations(10,14-17,31-36,78), but the samples mainly represented older individuals with limited transcriptional profiling performed. In recent years, the advent of single cell technologies has led to in-depth characterization of human pancreatic cancer and its accompanied tumor microenvironment(24,26,60,79,80). In those studies, tumor specific changes are assessed in comparison with adjacent “normal” pancreas which is often desmoplastic and inflamed. Thus, the characteristics of the human pancreas, and specifically its complement of non-epithelial cells, have largely remained unexplored.

Organ donation provides a unique opportunity to collect human tissue in a controlled manner in which conditions can be optimized to suspend cellular function in a physiologic state, allowing for virtually no warm ischemic time. Histopathology analysis revealed that the majority of our cohort of donor pancreata (slightly over half) presented with PanIN, neoplastic lesions considered progenitors to pancreatic cancer. While a similar prevalence was observed before, it was in the context of autopsy studies with patients in the 7^th^ and 8^th^ decades of life(14-17,78). Other investigators have detected a high prevalence of PanINs in the pancreas, but their studies were based on tumor adjacent normal areas(36). In contrast, our study indicate that PanIN are common in the pancreas of individual with no known pancreas pathology and occur early in life. While there was a slight increase in prevalence in the elderly, the broad distribution of ages counters prior beliefs that PanINs accumulate over time. To ensure accurate assessment of PanIN burden, multiple samples were taken from the head, body and tail of each donor organ for histologic analysis. We observed no overt difference in prevalence by anatomic region; PanINs, when present, were typically multi-focal and widely dispersed throughout the gland, consistent with recent reports(36).

In mice, spontaneous PanIN is virtually never observed; the lack of PanIN in mice might reflect the age difference (mice rarely live to two years of age, and most studies are conducted on animals of less than a year old). Further, laboratory mice live in a highly controlled environment, with predictable schedules and diet and no risk factors such as environmental pollution and tobacco use. In addition to PanIN, we observed frequent areas of acinar ductal metaplasia (ADM), dedifferentiation to acinar cells to duct-like cells. In mouse, acinar cells are the prevalent source of PanIN, and PanIN is preceded by ADM(81). Duct cells can also give rise to pancreatic cancer but they appear to do so bypassing the PanIN stage(82,83). The origin of PanIN and pancreatic cancer is still under debate, and the presence of ADM in human samples has been questioned; analysis of donor organs showed common ADM and PanIN, but no clear continuality between the two; PanINs were more often contiguous with ducts, pointing to a ductal origin as in previous reports(84-86).

To better understand PanIN lesions in the context of a non-diseased pancreas, we performed multiparametric analysis of donor pancreatic tissue, pancreatic adenocarcinoma and adjacent uninvolved tissue. We determined that PanIN lesions and their surrounding microenvironment had a desmoplastic stroma which was vastly different from normal pancreatic acinar tissue including acinar to ductal metaplasia. PanIN stroma had a unique inflammatory infiltrate that included myeloid cells as well as T helper and cytotoxic T cells. In contrast to tumors, where regulatory T cells (Tregs) are prevalent(51), we detected few or no Tregs in PanINs. This is also in contrast to murine models (and possibly tumor associated PanINs) which harbor regulatory T cells adjacent to ADM and PanIN(51). Similar to pancreatic ducts, human PanIN lesions were surrounded an array of collagen and fibroblasts. While both expressed fibroblast activation protein (FAP), fibroblasts surrounding PanIN lesions also expressed vimentin and smooth muscle actin (SMA), the latter a marker of fibroblast activation. Whether this stroma’s main function is to restrain PanIN progression, as shown in experimental models(87,88) or whether it promotes PanIN progression remains to be determined.

We sought to understand the transcriptional signature of epithelial and stromal cells in the normal pancreas. Our analysis revealed heterogeneous populations of fibroblasts and other mesenchymal cells, such as endothelial cells and pericytes, as well as abundant myeloid cells. As these same cell populations are common in pancreatic cancer stroma, we compared their characteristics in the normal organ compared to tumors. Interestingly, non-epithelial cell populations were largely distinct based on their origin, with gene expression signatures depending on their origin from the normal pancreas or from tumors. Differences in gene expression were most notable in fibroblasts, with markers of cancer associated fibroblasts (CAFs), such as the recently described *LRRC15* (89) exclusively expressed in the latter. In contrast, myeloid cells were less distinct; further analysis revealed that myeloid cells in the normal pancreas displayed a gene signature reminiscent of tumor associated macrophages (TAMs); whether the TAM-like status was present originally or is a byproduct of time on life support is unfortunately impossible to determine. Fibroblasts in pancreatic cancer have been described as heterogeneous, with smooth muscle actin high, myofibroblast-like myCAFs existing in parallel with inflammatory fibroblasts (iCAFs) and antigen presenting CAFs (apCAFs)(23); additional heterogeneity is due to cell of origin and expression of other specific markers(50) Interestingly, in our cohort fibroblast populations showed a distinct shift in tumors compared to normal pancreas, the latter not clearly reflecting this CAF classification but showing a similar level of heterogeneity.

Lastly, we interrogated the transcriptional signature of PanINs from the healthy pancreas compares with normal acinar and ductal cells, tumor associated PanINs and tumor cells. Of note, tissue dissociation during processing of single cell RNA sequencing samples makes it impossible to identify PanIN cells within the samples, as spatial and histological information is lost. To bypass this limitation, we leveraged spatial transcriptomics, then integrated transcriptomic signatures from spatial data with single cell RNA sequencing data. Interestingly, we observed that PanINs in the normal pancreas and in the tumor transcriptionally have a high degree of concordance, although do have some distinct features. Additionally, PanINs are transcriptionally distinct from ducts, and are markedly different from ADM. Surprisingly, integration of PanIN signatures with our single cell data revealed that PanINs are transcriptionally closely related to malignant tumors. This finding raises multiple questions; key among those are why PanINs, found frequently in the population, rarely progress to invasive disease given the relatively low incidence of pancreatic cancer. The endeavor to identify mechanisms that prevent PanIN progression bears important clinical significance, as results may have the possibility to be leveraged therapeutically.

## Methods

### Donor Sample Procurement

Donor pancreata not eligible for transplant or for whom there were no eligible recipients were collected at the Gift of Life Michigan Donor Care Center. While maintaining blood flow, the superior mesenteric arteries and celiac arteries were cross clamped and pancreas was recovered and immediately placed in physiologic organ preservation solution on ice and transported to the University of Michigan, approximately 6 miles away. Acquisition of donor pancreata for research purposes was approved by the Gift of Life research review group. Acquisition of tumor samples was described previously(24) and approved by the University of Michigan Institutional Review Board (HUM00025339).

### Tissue processing

Upon arrival to the laboratory, the donor pancreas was dissected by a clinically-trained pancreatobiliary surgeon. The organ was dissected into head, body, and tail, with portions of each placed into DMEM media with 1%BSA/10μM Y27632 or 10% formalin for single cell sequencing or paraffin embedding, respectively. For single cell processing, tissue was minced into 1mm^3^ pieces, then digested with 1 mg/mL collagenase P for 20-30 min at 37°C with gentle agitation. Digested tissue was rinsed three times with DMEM/1%BSA/10μM Y27632, then filtered through a 40μm mesh. Resulting cells were submitted to the University of Michigan Advanced Genomics Core for single cell sequencing using the 10x Genomics Platform.

### Multiplex Fluorescent Immunohistochemistry (mfIHC)

Formalin-fixed, paraffin embedded (FFPE) tissue slides were rehydrated in triplicate with xylene, followed by a single submersion in 100% ethanol, 95% ethanol, and 70% ethanol, respectively. Rehydrated slides are then washed in neutral buffered formalin and rinsed with deionized water. Antigen retrieval was performed with sodium citrate, pH 6.0 buffer for membrane and cytoplasmic epitopes and tris-EDTA, pH 9.0 buffer for nuclear epitopes. Tissue was blocked using endogenous peroxidase at room temperature, followed by 10% donkey serum overnight at 4°C. Primary antibodies were diluted in 5% donkey serum in TBST and incubated overnight at 4°C. Subsequent multiplex staining was completed using a 1% BSA block. Upon completion of the multiplex, tissue samples were rinsed in deionized water and mounted with DAPI ProLong^™^ Diamond Antifade Mountant (Thermo Fisher Scientific). Images were taken using the Vectra® Polaris™ Work Station (Akoya Biosciences). Antibodies/dilutions used are listed in Supplementary Table 4.

### Multiplex Immunofluoresence Quantification

Images were acquired using the Mantra^™^ Quantitative Pathology Work Station (Akoya Biosciences). A minimum of ten images were acquired from each tissue seciton. All cube filters were used for each image capture (DAPI, CY3, CY5, CY7, Texas Red, Qdot) and the saturation protection feature was utilized. After all images were acquired, images were analyzed using inForm® Cell Analysis^™^ software (Akoya Biosciences).

Using this software, acinar, ductal, acinar to ductal metaplasia and PanIN samples were batch analyzed by their separate diagnoses which was confirmed by a pathologist. Cell segmentation was completed using DAPI as a basis of cell location and nuclear size and all cells segmented into the following subsets (nucleus, cytoplasm, and membrane). Using the automated training software, basic phenotypes were created. For the fibroblast panel, this included Vimentin^+^, smooth muscle actin (SMA^+^) and PDGFR^+^. For the immune based panel, this included CD3^+^, CD8^+^, CD163^+^, PanCK^+^ and FoxP3^+^.

Software output consisting of mean fluorescent intensity (mfi) of each antibody-fluorophore pair, basic phenotypes, and x and y coordinates were acquired for further processing to determine relative population of each cell type.

### RNA scope Multiplex Fluorescent Detection with Immunofluorescence

RNA scope Multiplex Fluorescent Detection was performed according to instructions provided by the manufacture (Advance Cell Diagnostics; ACD). Briefly, 5µm thick formalin fixed paraffin embedded normal pancreas tissue sections were mounted on charged slides and baked at 60°C for 1 hour, deparaffinized, dehydrated, and washed with 0.1% Tween-20 RNAse-free 1x phosphate-buffered saline (PBST) three times. This was followed by incubation with hydrogen peroxide for 10 minutes at room temperature and target retrieval for 15 minutes at 98°C. Slides were then blocked with Co-Detection antibody diluent (ACD) for 30 minutes prior to incubation with Claudin 18 primary antibody (1:100 in co-detection antibody diluent) for 24 hours at 4°C. The next day the tissue sections were fixed with formalin and treated with ProteasePlus Reagent (Advanced Cell Diagnostics) for 13 minutes at 40°C. Amplification and signal enhancement (AMP) were performed for two different probes/channels (C1, C2). The TFF1-C1 probe was 1x concentration, while the CCL2-C2 probe was at 50x concentration. The probes were diluted as per manufacturer’s instructions and slides were incubated with them at 40°C for 2 hours. Then the slides were washed with RNA scope washing buffer (ACD) twice. Signal for each of the probes was amplified with AMP reagents, horseradish peroxidase, and tyramide signal amplification kit (ACD) at 40°C. Slides were then washed with PBST (PBS with 0.1% Tween20) three times, incubated with anti-rabbit secondary Alexa Fluor IgG (H + L) antibody (1:400 in co-detection antibody diluent) for 1 hour at room temperature. Finally, slides were incubated with DAPI for 15 minutes and washed three times with PBST prior to being mounted with ProLong Diamond Antifade.

For details on Immunofluorescence Assessment with Tyramide Signal Amplification, please refer to Supplementary Methods.

### Single-cell RNA sequencing/Pseudobulk DE/Differential Abundance/Geneset Scoring/Spatial Transcriptomics Data Analysis

Please refer to Supplementary Methods for full details:

## Acknowledgements

We sincerely thank Gift of Life Michigan and donors and their families who have generously provided organs for this research. We thank T. Tamsen, M. Hogan and J. Opp from the University of Michigan Advanced Genomics Core for their assistance with single cell RNA sequencing and spatial transcriptomics.

Work in the M.P.d.M. laboratory was supported by NIH/NCI grants R01-CA268426, R01-CA260752, R01-CA271510, R01-CA264843, U01-CA224145, U01-CA274154, and U54CA274371. Work in the T.L.F laboratory was supported by NIH/NCI grant U01CA274154, NIH NIDDK 5R01DK128102, and VA BLR&D Merit Award 5I01BX005777.

E.S.C. was supported by the VA BLR&D CDA IK2BX005875, the American College of Gastroenterology CDA, and by NIH NIDDK T32-DK094775. A.M.E was supported by the Rackham International Student Fellowship. P.K. was supported by NIH/NIAID T32-AI007413 and NIH/NIGMS T32-GM113900. J.L was supported by NIH/NIGMS T32GM007863. K.D. was supported by NIH/NCI F31-CA265085-01A1. AR was supported by NCI Grants R37-CA214955, CCSG Bioinformatics Shared Resource 5 P30 CA046592. F.B. was funded by the Association for Academic Surgery Joel J. Roslyn Award and NCI R01-CA271510. J.S. was funded by NIH NCI K08CA234222 and R37CA262209. The funders had no role in study design, data collection and analysis, decision to publish or preparation of the manuscript.

## Table Legends

Supplemental Table 1: Demographics and clinical data for donor Gife of Life Samples

Supplemental Table 2: Putative Ligand-Receptor pairs comparing all cell types from single cell sequencing of tumor samples compared to healthy samples.

Supplemental Table 3: Differential gene expression using linear mixed model analysis of cell types from spatial transcriptomic ROIs.

Supplemental Table 4: Antibodies used for multiplex immunofluoresence. Supplemental Table 5: Antibodies and RNAScope probe used for co-IF/ISH.

## Supplemental Figure Legends

Supplemental Fig 1

A) Representative slide scans from head (left), body (middle), and tail (right) of tissue sections from a donor pancreas.

B) Representative H&E sections of acinar, ADM (acinar-to-ductal metaplasia), normal duct, and PanIN structures in a donor pancreas.

C) H&E from a DCD (donation after cardiac death) organ, showing extensive tissue degradation.

Supplemental Fig 2

A) mfIHC composite images of formalin-fixed, paraffin-embedded donor tissue specimens (each row taken from different donor organs), highlighting acinar, normal duct, ADM (acinar-to-ductal), and PanIN structures. Antibodies and colors of immune panel used in the legend below.

B) mfIHC composite images of formalin-fixed, paraffin-embedded donor tissue specimens (each row taken from different donor organs), highlighting acinar, normal duct, ADM (acinar-to-ductal), and PanIN structures. Antibodies and colors of fibroblast panel used in the legend below.

Supplemental Fig 3

A) Dotplot showing average expression and percent cells of top expressing markers in each cell cluster.

B) Dotplot showing markers defining different myeloid populations.

C) Single-cell-resolution heatmap of top expressed genes in each myeloid cell population identified.

Supplemental Fig 4

A) Dotplot showing markers defining different lymphocyte populations.

B) Single-cell-resolution heatmap of top expressed genes in each T&NK cell population identified.

C) Single-cell-resolution heatmap of top expressed genes in each B cell and plasma cell population identified.

Supplemental Fig 5

A) Dotplot showing markers defining different populations endothelial, fibroblast, and pericyte populations.

B) Featureplots of select fibroblast and endothelial markers on UMAP of extracted fibroblast, endothelial, and pericyte cells.

C) Single-cell-resolution heatmap of top expressed genes in each fibroblast and pericyte population.

D) Single-cell-resolution heatmap of top expressed genes in each endothelial cell population.

Supplemental Fig 6

A) Histogram of cell type abundance of all cell populations by sample. Blue samples represent healthy samples, magenta samples represent adjacent normal samples, and orange samples represent tumor samples.

B) UMAP overlay of disease states on all cells from single cell dataset of healthy, adjacent normal, and tumor samples.

C) Heatmap of the differnetial interaction strength comparing tumor samples to healthy samples. Y-axis represents outgoing signaling sent from cell populations while x-axis represents incoming signaling received to cell populations.

D) (left) Circle interaction plot showing differentially outgoing signaling strength upregulated in tumors compared to healthy in fibroblasts, endothelial cells, and pericytes. (right) Circle interaction plot showing differentially outgoing signaling strength upregulated in tumors compared to healthy in all cell types. Red lines indicated increased signaling in tumors, blue lines indicate decreased signaling in tumors.

Supplemental Fig 7

A) Histogram of cell type abundance of myeloid populations by sample. Blue samples represent healthy samples, magenta samples represent adjacent normal samples, and orange samples represent tumor samples.

B) Dotplot showing markers defining myeloid populations.

C) Treeplot of gene ontology functional annotation on pseudobulk aggregated tumor myeloid cells vs. healthy myeloid cells. Dot size represents number of genes in each pathway, and color bar represents adjusted p-value.

D) Single-cell-resolution heatmap of top expressed genes in fibroblast/pericyte populations in merged healthy and tumor samples.

E) Featureplots of select fibroblast markers on UMAP of extracted fibroblast and pericyte cells from merged healthy and tumor samples.

Supplemental Fig 8

A) (left) UMAP of extracted NK and T cell s from single cell dataset of healthy, adjacent normal, and tumor samples. Populations are identified by color. (right) UMAP overlay of disease states on extracted fibroblast/pericyte cells from single cell dataset of healthy, adjacent normal, and tumor samples.

B) Dotplot showing markers defining NK and T populations.

C) PCA plots of pseudobulk-aggregated counts from all cells, myeloid cells, t-cells, and fibroblasts. Each dot represents one aggregated single cell sequencing sample.

D) Treeplot of gene ontology functional annotation on pseudobulk aggregated tumor T cells vs. healthy T cells. Dot size represents number of genes in each pathway, and color bar represents adjusted p-value.

Supplemental Fig 9

A) UMAP of extracted epithelial cells from single cell dataset of healthy, adjacent normal, and tumor samples. Numbers represent unbiased clustering.

B) Dotplot of top expressed markers in each of the epithelial clusters.

C) UMAP split by disease states on extracted epithelial cells from single cell dataset of healthy, adjacent normal, and tumor samples.

D) Single cell resolution heatmap of top expressed markers in each unbiased epithelial cluster.

E) AUCell geneset scoring mapped to epithelial single cell dataset of healthy, adjacent normal, and tumor samples using previously reported(60) acinar, ductal, panIN, and tumor gene signatures.

F) *KRAS* activity score mapped to epithelial single cell dataset of healthy, adjacent normal, and tumor samples utilizing previously reported(90) hallmark genesets.

Supplemental Fig 10

A) PCA plot of all segmented ROIs (each dot represents one ROI) pre- and post-batch correction.

B) Scheme for workflow of spatial trancriptomics analysis.

C) Violin plot of MUC5AC expression in spatial ROIs by cell type and disease state.

D) Heatmap of geneset enrichiment score using GSVA of Exocrine_like, ADEX, Classical, and Basal subyptes.

Supplemental Fig 11-14

Immunofluresence staining of serial sections of normal duct (ND), ADM, and PanIN lesions found in donor tissue. In order from top row to bottom row: Merged co-IF of MUC5AC (yellow) and MUC5B (blue). Single channel of MUC5AC staining. Single channel of MUC5B staining. Merge of co-IF of AQP1 (grey) and AQP3 (purple). Single channel of AQP1 staining. Single channel of AQP3 staining. Merge of co-IF of p-ERK (green) and CLDN18 (orange). Single channel of p-ERK staining. Single channel of CLDN18 staining.

## Notes

**Conflict of interest disclosure statement** The authors declare no competing interests for this manuscript.

### Competing Interest Statement

The authors have declared no competing interest.

