## Supplementary Figures for "Analysis of donor pancreata defines the transcriptomic signature and microenvironment of early pre-neoplastic pancreatic lesions"

Supp Fig 1

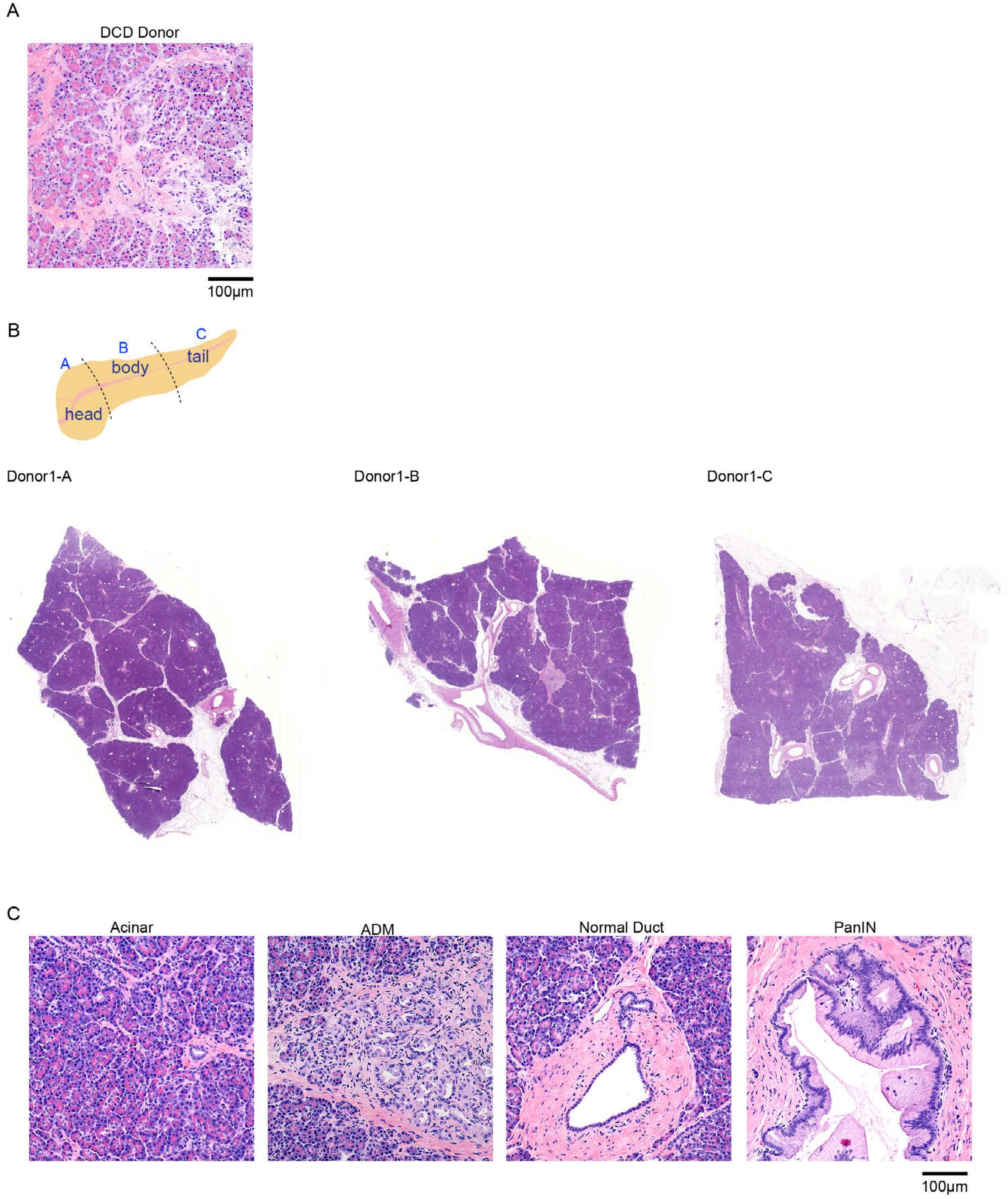
A
DCD Donor
100µm
B
A
B
C
head
body
tail
Donor1-A
Donor1-B
Donor1-C
C
Acinar
ADM
Normal Duct
PanIN
100µm

Supp Fig 2

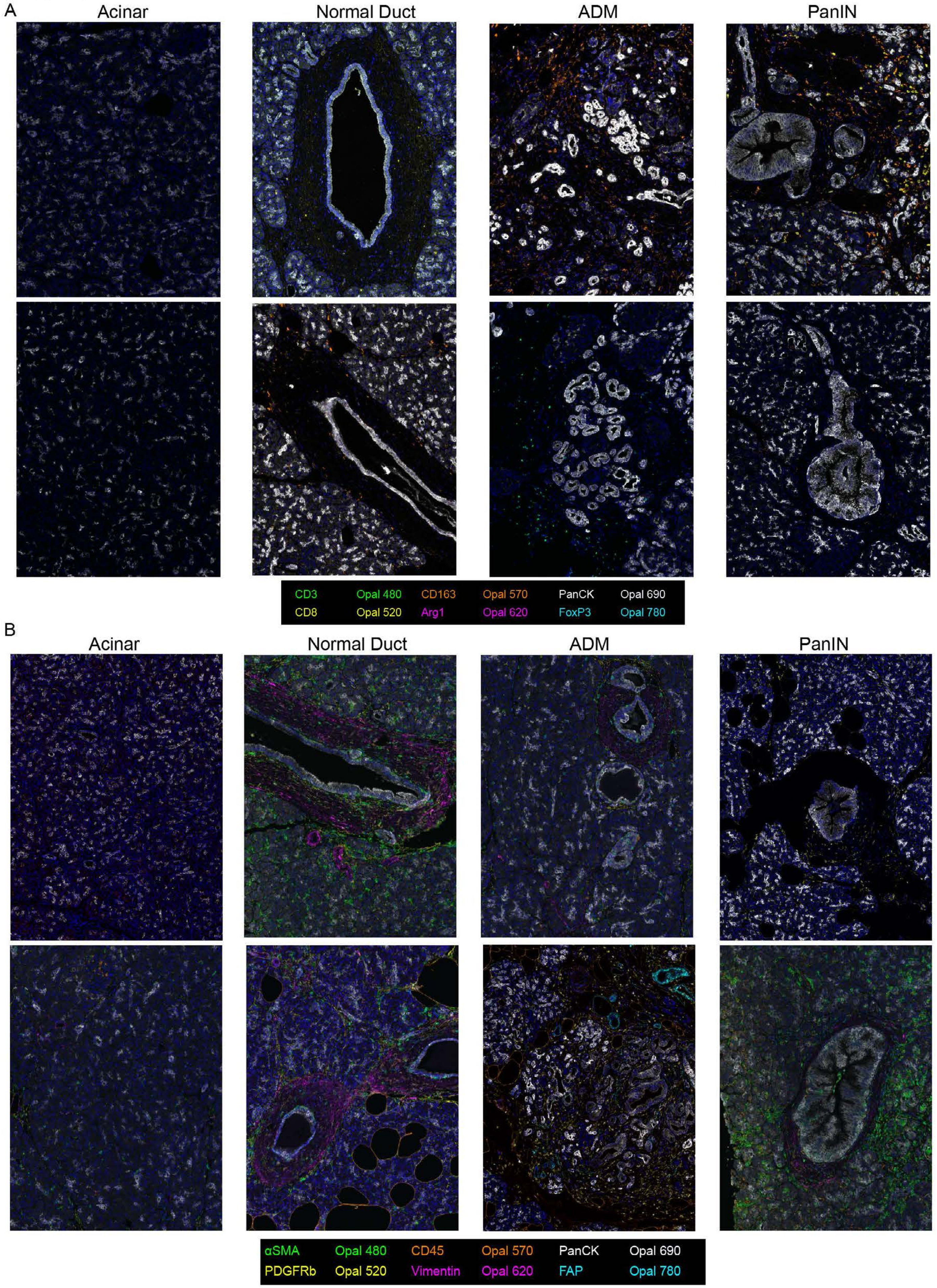

A
Acinar
Normal Duct
ADM
PanIN
CD3
Opal 480
CD163
Opal 570
PanCK
Opal 690
CD8
Opal 520
Arg1
Opal 620
FoxP3
Opal 780
B
Acinar
Normal Duct
ADM
PanIN
αSMA
Opal 480
CD45
Opal 570
PanCK
Opal 690
PDGFRb
Opal 520
Vimentin
Opal 620
FAP
Opal 780

Supp Fig 3

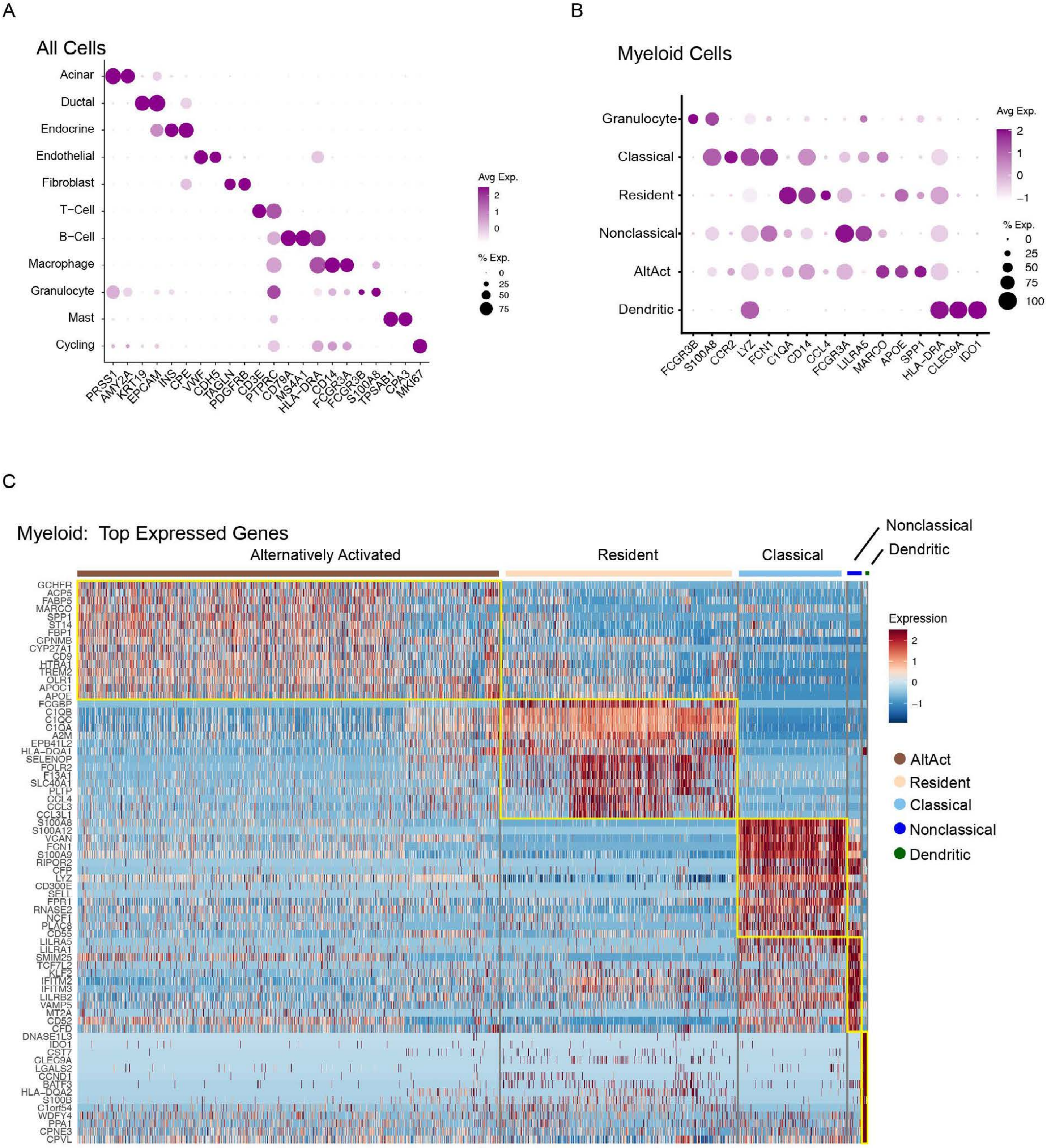
A
All Cells
Acinar
Ductal
Endocrine
Endothelial
Fibroblast
T−Cell
B−Cell
Macrophage
Granulocyte
Mast
Cycling
PRSS1 AMY2A KRT19 EPCAM INS CPE VWF CDH5 TAGLN PDGFRB CD3E PTPRC CD79A MS4A1 HLA−DRA CD14 FCGR3A FCGR3B S100A8 TPSAB1 CPA3 MKI67
Avg Exp.
2
1
0
% Exp.
0
25
50
75
B
Myeloid Cells
Granulocyte
Classical
Resident
Nonclassical
AltAct
Dendritic
FCGR3B S100A8 CCR2 LYZ FCN1 C1QA CD14 CCL4 FCGR3A LILRA5 MARCO APOE SPP1 HLA−DRA CLEC9A IDO1
Avg Exp.
2
1
0
−1
% Exp.
0
25
50
75
100
C
Myeloid: Top Expressed Genes
Alternatively Activated
Resident
Classical
Nonclassical
Dendritic
GCHFR ACP5 FABP5 MARCO SPP1 ST14 FBP1 GPNMB CYP27A1 CD9 HTRA1 TREM2 OLR1 APOC1 APOE FCGBP C1QB C1QC C1QA A2M EPB41L2 HLA−DQA1 SELENOP FOLR2 F13A1 SLC40A1 PLTP CCL4 CCL3 CCL3L1 S100A8 S100A12 VCAN FCN1 S100A9 RIPOR2 CFP LYZ CD300E SELL FPR1 RNASE2 NCF1 PLAC8 CD55 LILRA5 LILRA1 SMIM25 TCF7L2 KLF2 IFITM2 IFITM3 LILRB2 VAMP5 MT2A CD52 CFD DNASE1L3 IDO1 CST7 CLEC9A LGALS2 CCND1 BATF3 HLA−DQA2 S100B C1orf54 WDFY4 PPA1 CPNE3 CPVL
Expression
2
1
0
−1
AltAct
Resident
Classical
Nonclassical
Dendritic

Supp Fig 4

A

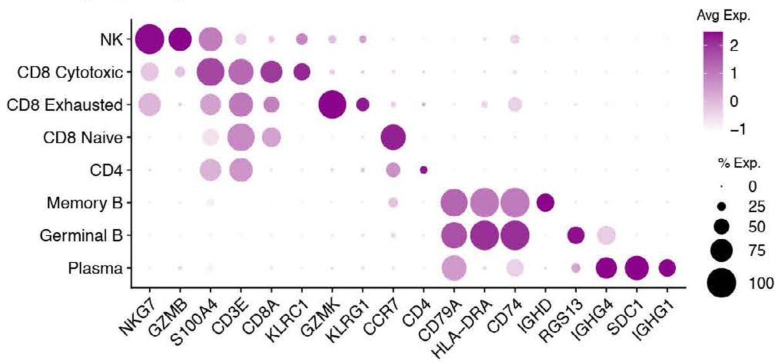

B

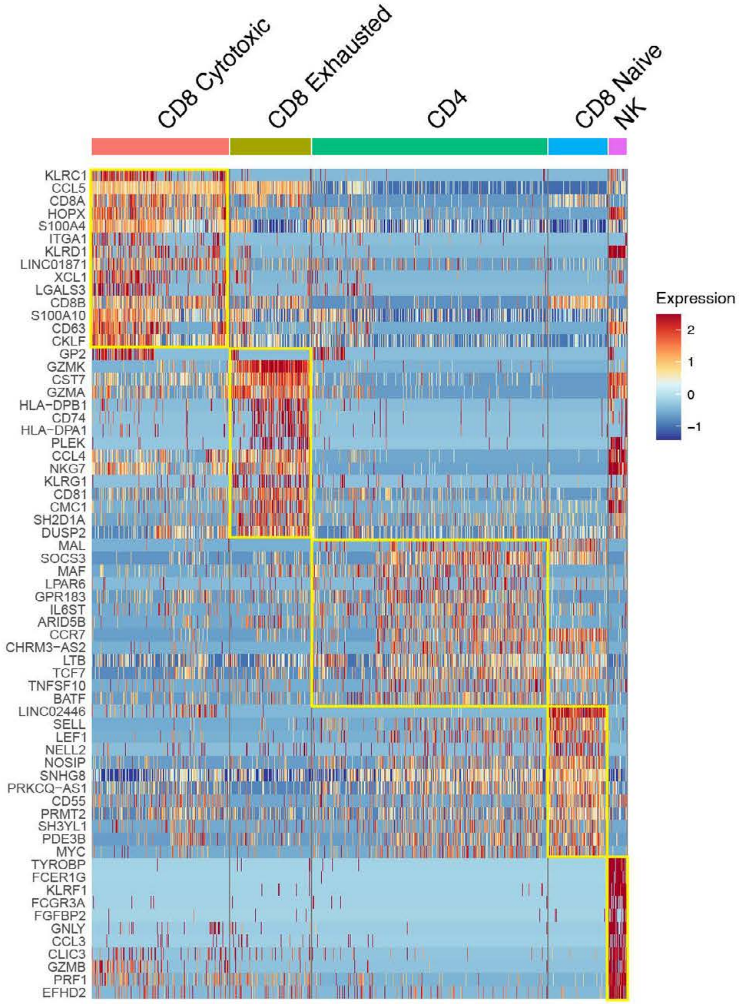

C

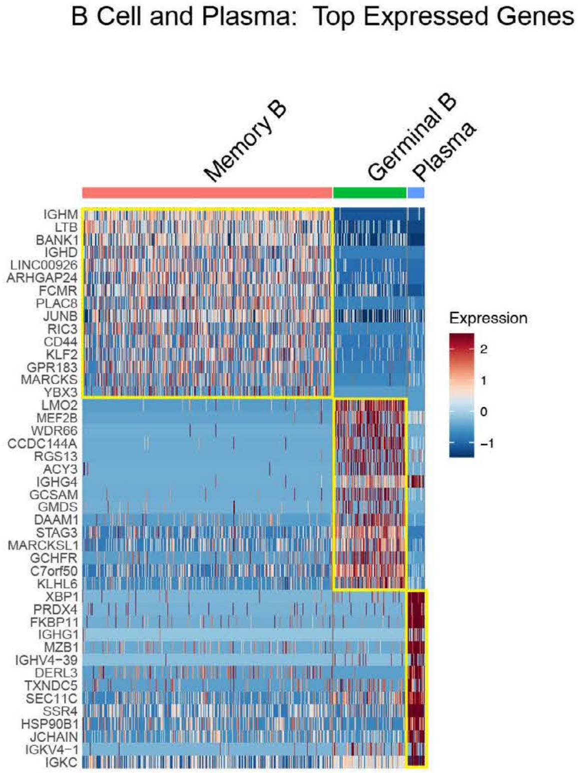

Supp Fig 5

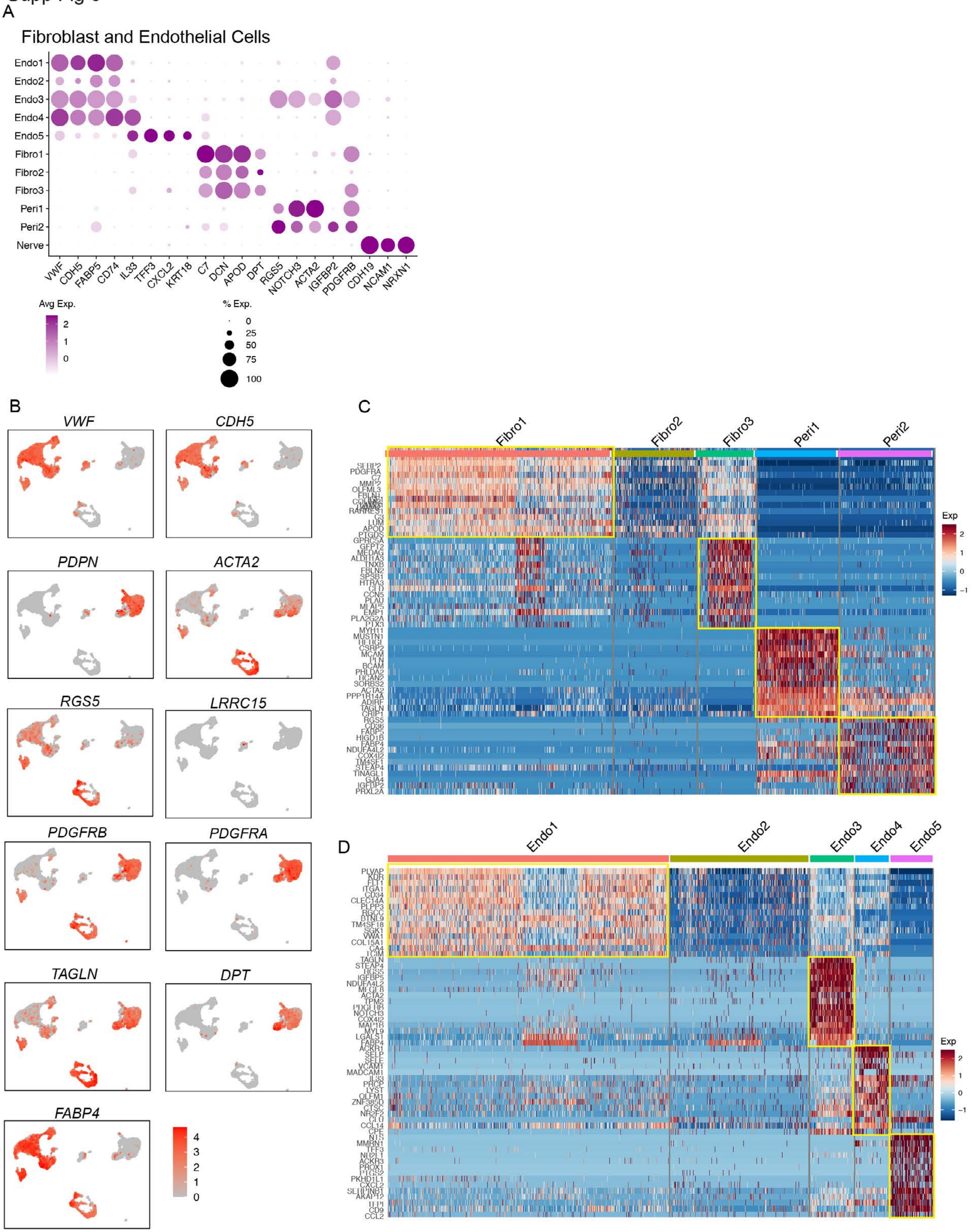
A
Fibroblast and Endothelial Cells
Endo1
Endo2
Endo3
Endo4
Endo5
Fibro1
Fibro2
Fibro3
Peri1
Peri2
Nerve
VWF CDH5 FABP5 CD74 IL33 TFF3 CXCL2 KRT18 C7 DCN APOD DPT RGS5 NOTCH3 ACTA2 IGFBP2 PDGFRB CDH19 NCAM1 NRXN1
Avg Exp.
2
1
0
% Exp.
0
25
50
75
100
B
VWF
CDH5
PDPN
ACTA2
RGS5
LRRC15
PDGFRB
PDGFRA
TAGLN
DPT
FABP4
4
3
2
1
0
C
Fibro1
Fibro2
Fibro3
Peri1
Peri2
Exp
2
1
0
−1
D
Endo1
Endo2
Endo3
Endo4
Endo5
Exp
2
1
0
−1

Supp Fig 6

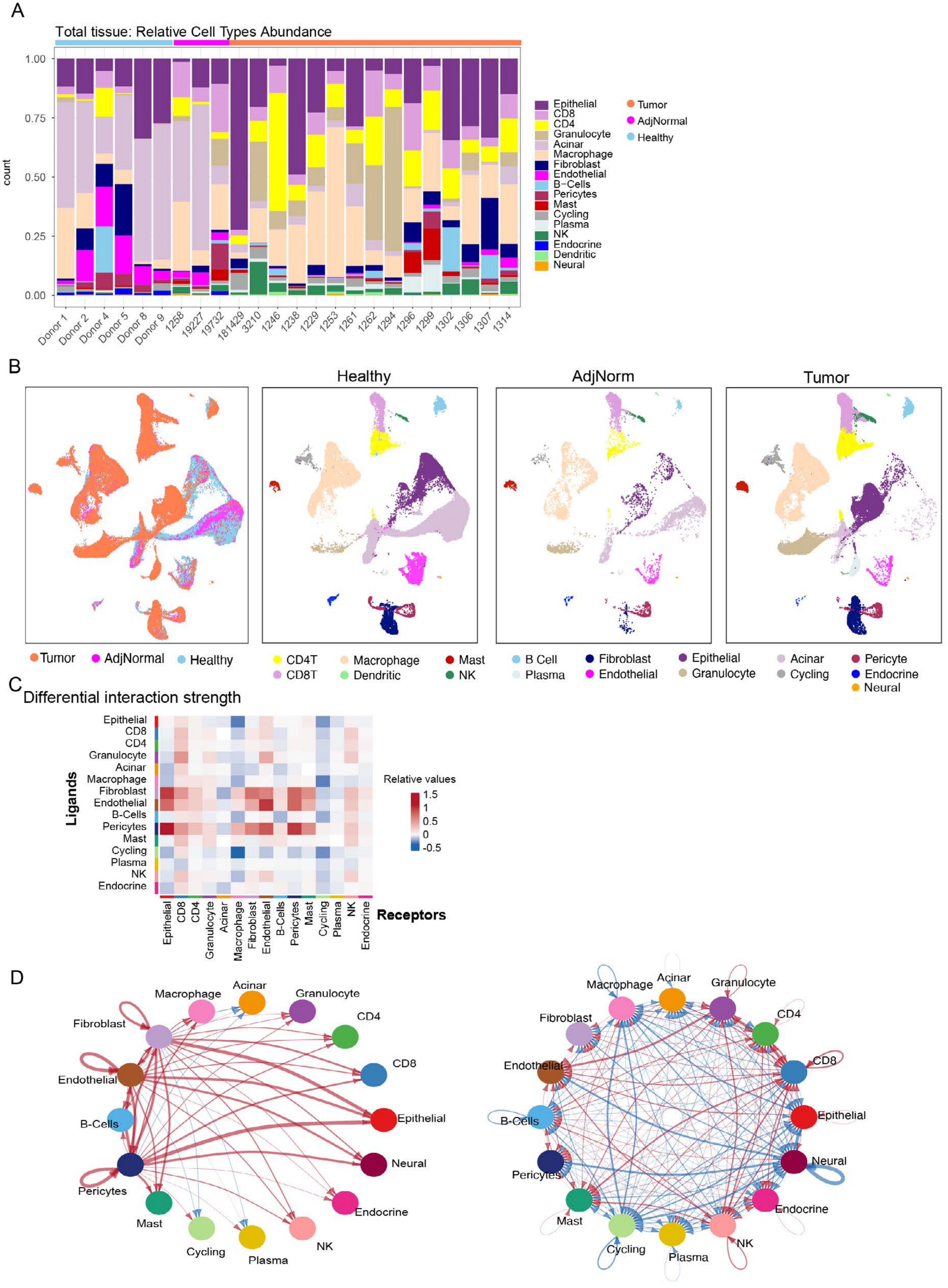

A
Total tissue: Relative Cell Types Abundance
count
1.00
0.75
0.50
0.25
0.00
Donor 1
Donor 2
Donor 4
Donor 5
Donor 8
Donor 9
1258
19227
19732
181429
3210
1246
1238
1229
1253
1261
1262
1294
1296
1299
1302
1306
1307
1314
Epithelial
CD8
CD4
Granulocyte
Acinar
Macrophage
Fibroblast
Endothelial
B−Cells
Pericytes
Mast
Cycling
Plasma
NK
Endocrine
Dendritic
Neural
Tumor
AdjNormal
Healthy
B
Healthy
AdjNorm
Tumor
Tumor
AdjNormal
Healthy
CD4T
CD8T
Macrophage
Dendritic
Mast
NK
B Cell
Plasma
Fibroblast
Endothelial
Epithelial
Granulocyte
Acinar
Cycling
Pericyte
Endocrine
Neural
C
Differential interaction strength
Ligands
Receptors
Relative values
1.5
1
0.5
0
-0.5
Epithelial
CD8
CD4
Granulocyte
Acinar
Macrophage
Fibroblast
Endothelial
B-Cells
Pericytes
Mast
Cycling
Plasma
NK
Endocrine
D
Macrophage
Acinar
Granulocyte
CD4
CD8
Epithelial
Neural
Endocrine
NK
Plasma
Cycling
Mast
Pericytes
B-Cells
Endothelial
Fibroblast

Supp Fig 7

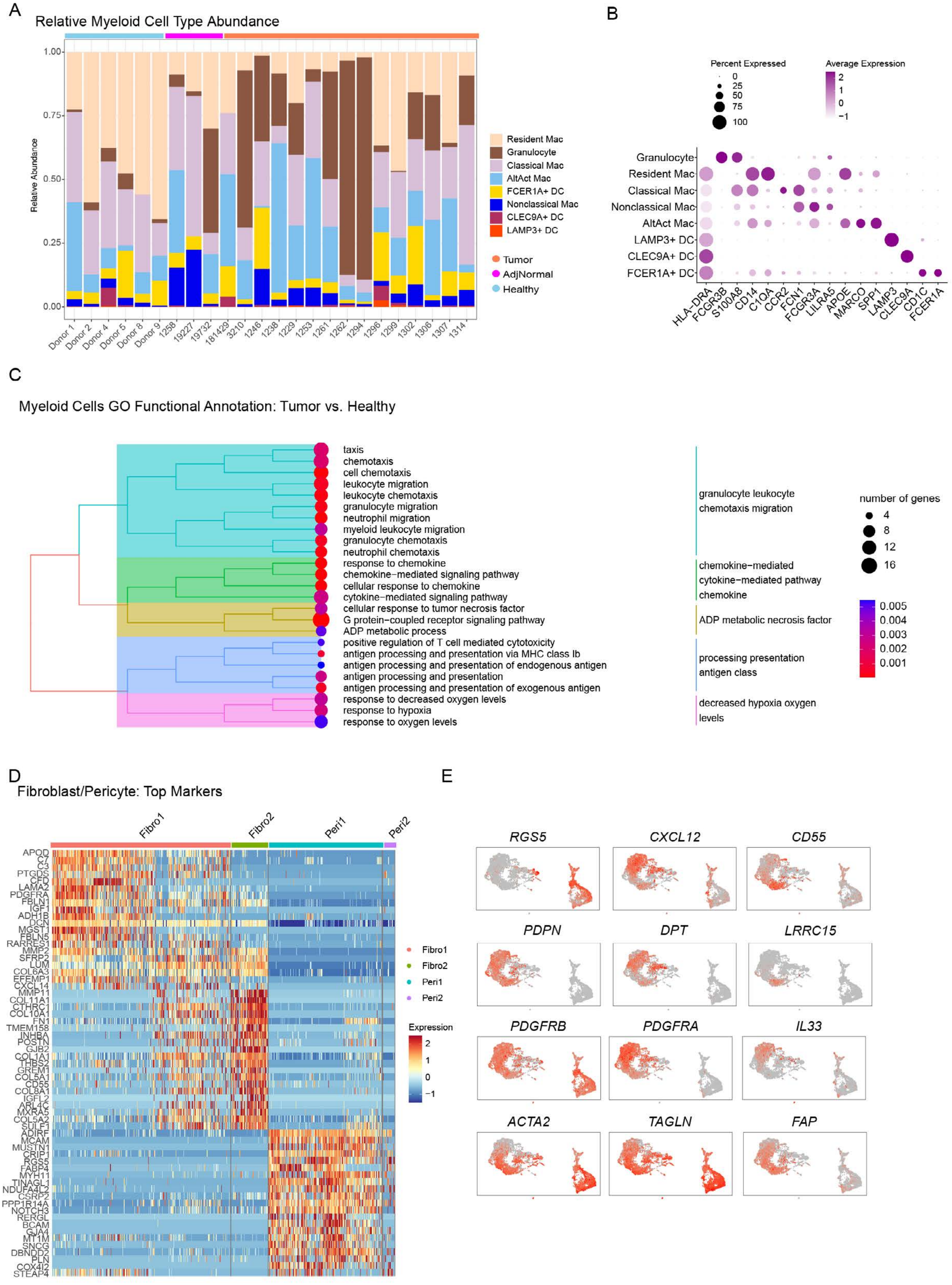
A
Relative Myeloid Cell Type Abundance
Relative Abundance
1.00
0.75
0.50
0.25
0.00
Donor 1
Donor 2
Donor 4
Donor 5
Donor 8
Donor 9
1258
19227
19732
181429
3210
1246
1238
1229
1253
1261
1262
1294
1296
1299
1302
1306
1307
1314
Resident Mac
Granulocyte
Classical Mac
AltAct Mac
FCER1A+ DC
Nonclassical Mac
CLEC9A+ DC
LAMP3+ DC
Tumor
AdjNormal
Healthy
B
Percent Expressed
0
25
50
75
100
Average Expression
2
1
0
−1
Granulocyte
Resident Mac
Classical Mac
Nonclassical Mac
AltAct Mac
LAMP3+ DC
CLEC9A+ DC
FCER1A+ DC
HLA−DRA
FCGR3B
S100A8
CD14
C1QA
CCR2
FCN1
FCGR3A
LILRA5
APOE
MARCO
SPP1
LAMP3
CLEC9A
CD1C
FCER1A
C
Myeloid Cells GO Functional Annotation: Tumor vs. Healthy
taxis
chemotaxis
cell chemotaxis
leukocyte migration
leukocyte chemotaxis
granulocyte migration
neutrophil migration
myeloid leukocyte migration
granulocyte chemotaxis
neutrophil chemotaxis
response to chemokine
chemokine−mediated signaling pathway
cellular response to chemokine
cytokine−mediated signaling pathway
cellular response to tumor necrosis factor
G protein−coupled receptor signaling pathway
ADP metabolic process
positive regulation of T cell mediated cytotoxicity
antigen processing and presentation via MHC class Ib
antigen processing and presentation of endogenous antigen
antigen processing and presentation
antigen processing and presentation of exogenous antigen
response to decreased oxygen levels
response to hypoxia
response to oxygen levels
granulocyte leukocyte chemotaxis migration
chemokine−mediated cytokine−mediated pathway chemokine
ADP metabolic necrosis factor
processing presentation antigen class
decreased hypoxia oxygen levels
number of genes
4
8
12
16
0.005
0.004
0.003
0.002
0.001
D
Fibroblast/Pericyte: Top Markers
Fibro1
Fibro2
Peri1
Peri2
APOD
C7
C3
PTGDS
CFD
LAMA2
PDGFRA
FBLN1
IGF1
ADH1B
DCN
MGST1
FBLN5
RARRES1
MMP2
SFRP2
LUM
COL6A3
EFEMP1
CXCL14
MMP11
COL11A1
CTHRC1
COL10A1
FN1
TMEM158
INHBA
POSTN
GJB2
COL1A1
THBS2
GREM1
COL5A1
CD55
COL8A1
IGFL2
ARL4C
MXRA5
COL5A2
SULF1
ADIRF
MCAM
MUSTN1
CRIP1
RGS5
FABP4
MYH11
TINAGL1
NDUFA4L2
CSRP2
PPP1R14A
NOTCH3
RERGL
BCAM
GJA4
MT1M
SNCG
DBNDD2
PLN
COX4I2
STEAP4
Fibro1
Fibro2
Peri1
Peri2
Expression
2
1
0
−1
E
RGS5
CXCL12
CD55
PDPN
DPT
LRRC15
PDGFRB
PDGFRA
IL33
ACTA2
TAGLN
FAP

Supp Fig 8

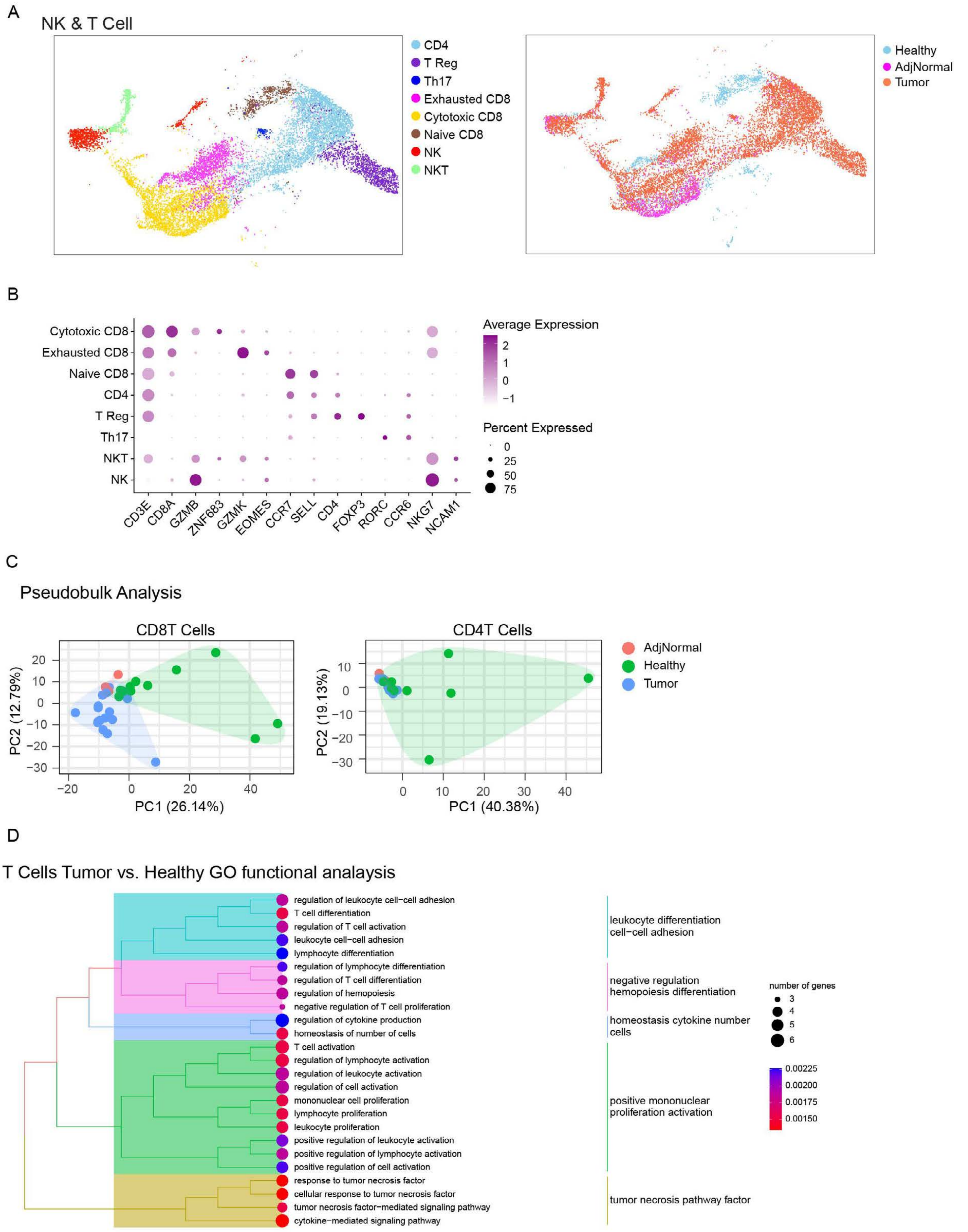
A
NK & T Cell
CD4
T Reg
Th17
Exhausted CD8
Cytotoxic CD8
Naive CD8
NK
NKT
Healthy
AdjNormal
Tumor
B
Cytotoxic CD8
Exhausted CD8
Naive CD8
CD4
T Reg
Th17
NKT
NK
CD3E
CD8A
GZMB
ZNF683
GZMK
EOMES
CCR7
SELL
CD4
FOXP3
RORC
CCR6
NKG7
NCAM1
Average Expression
2
1
0
−1
Percent Expressed
0
25
50
75
C
Pseudobulk Analysis
CD8T Cells
PC2 (12.79%)
PC1 (26.14%)
CD4T Cells
PC2 (19.13%)
PC1 (40.38%)
AdjNormal
Healthy
Tumor
D
T Cells Tumor vs. Healthy GO functional analaysis
regulation of leukocyte cell−cell adhesion
T cell differentiation
regulation of T cell activation
leukocyte cell−cell adhesion
lymphocyte differentiation
regulation of lymphocyte differentiation
regulation of T cell differentiation
regulation of hemopoiesis
negative regulation of T cell proliferation
regulation of cytokine production
homeostasis of number of cells
T cell activation
regulation of lymphocyte activation
regulation of leukocyte activation
regulation of cell activation
mononuclear cell proliferation
lymphocyte proliferation
leukocyte proliferation
positive regulation of leukocyte activation
positive regulation of lymphocyte activation
positive regulation of cell activation
response to tumor necrosis factor
cellular response to tumor necrosis factor
tumor necrosis factor−mediated signaling pathway
cytokine−mediated signaling pathway
leukocyte differentiation cell−cell adhesion
negative regulation hemopoiesis differentiation
homeostasis cytokine number cells
positive mononuclear proliferation activation
tumor necrosis pathway factor
number of genes
3
4
5
6
0.00225
0.00200
0.00175
0.00150

Supp Fig 9

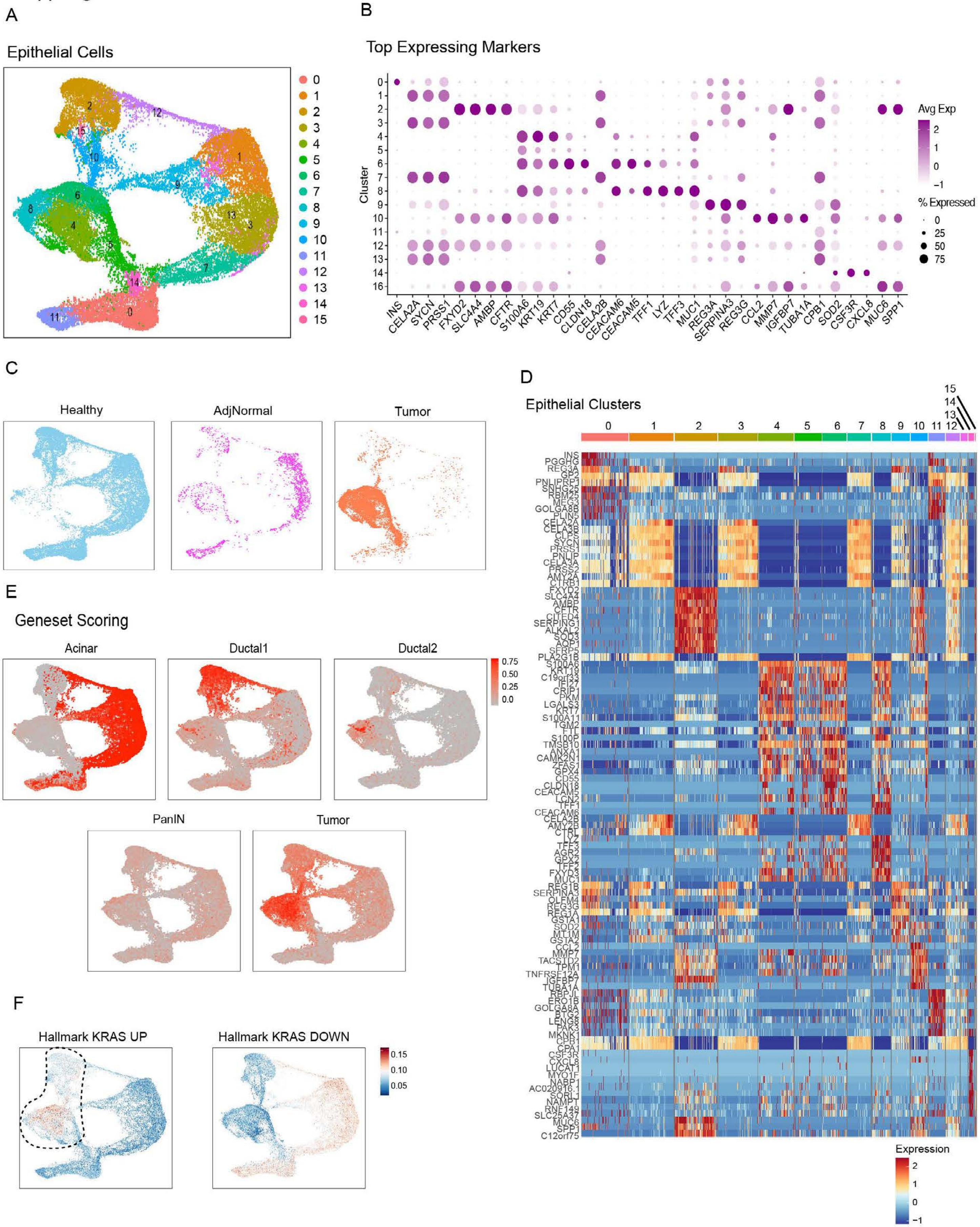

A
Epithelial Cells
B
Top Expressing Markers
Cluster
INS
CELA2A
SYCN
PRSS1
FXYD2
SLC4A4
AMBP
CFTR
S100A6
KRT19
KRT7
CD55
CLDN18
CELA2B
CEACAM6
CEACAM5
TFF1
LYZ
TFF3
MUC1
REG3A
SERPINA3
REG3G
CCL2
MMP7
IGFBP7
TUBA1A
CPB1
SOD2
CSF3R
CXCL8
MUC6
SPP1
Avg Exp
% Expressed
C
Healthy
AdjNormal
Tumor
D
Epithelial Clusters
Expression
E
Geneset Scoring
Acinar
Ductal1
Ductal2
PanIN
Tumor
F
Hallmark KRAS UP
Hallmark KRAS DOWN

Supp Fig 10

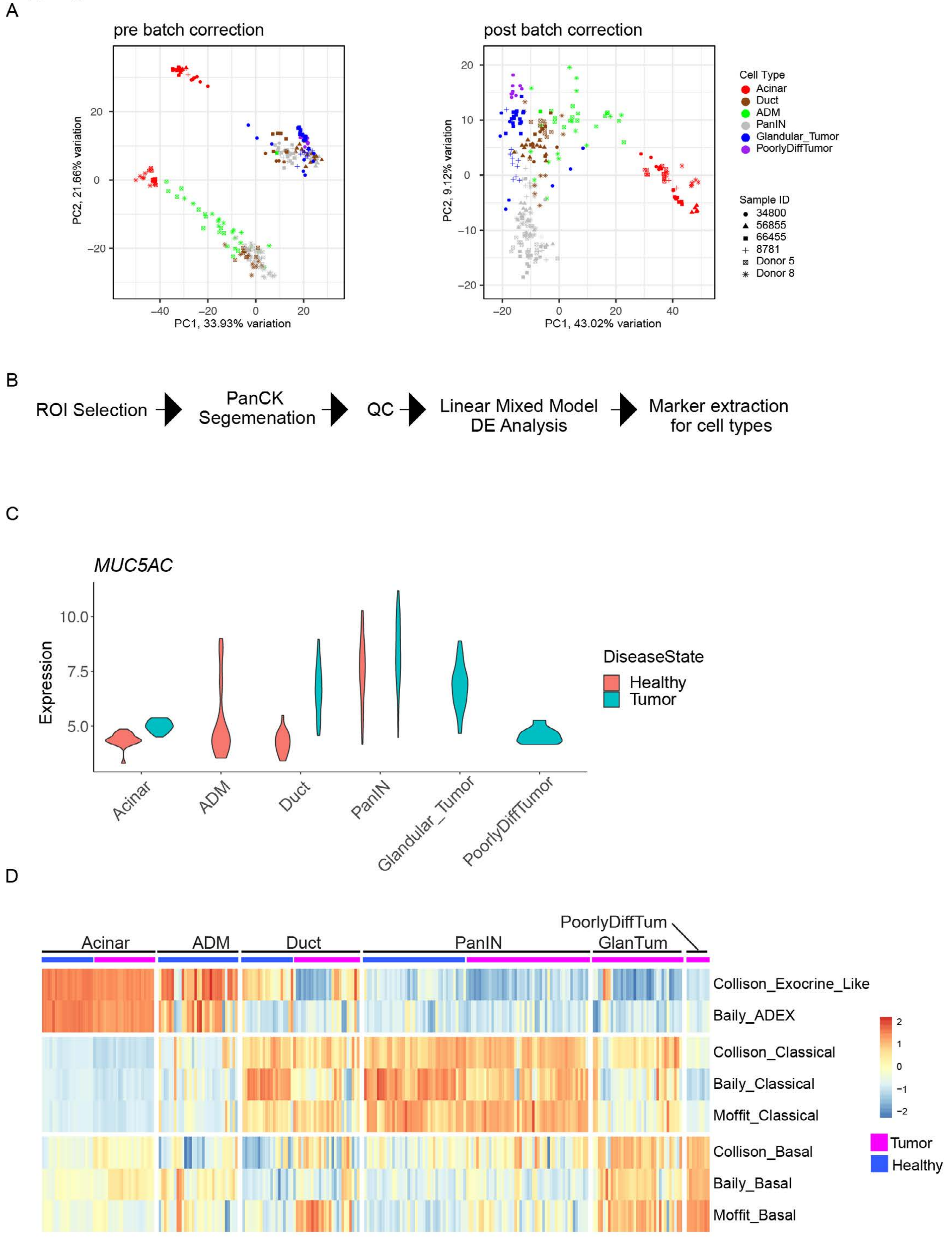
A
pre batch correction
PC2, 21.66% variation
PC1, 33.93% variation
post batch correction
PC2, 9.12% variation
PC1, 43.02% variation
Cell Type
Acinar
Duct
ADM
PanIN
Glandular_Tumor
PoorlyDiffTumor
Sample ID
34800
56855
66455
8781
Donor 5
Donor 8
B
ROI Selection
PanCK Segemenation
QC
Linear Mixed Model DE Analysis
Marker extraction for cell types
C
MUC5AC
Expression
DiseaseState
Healthy
Tumor
Acinar
ADM
Duct
PanIN
Glandular_Tumor
PoorlyDiffTumor
D
Acinar
ADM
Duct
PanIN
GlanTum
PoorlyDiffTum
Collison_Exocrine_Like
Baily_ADEX
Collison_Classical
Baily_Classical
Moffit_Classical
Collison_Basal
Baily_Basal
Moffit_Basal
Tumor
Healthy

Supp Fig 11

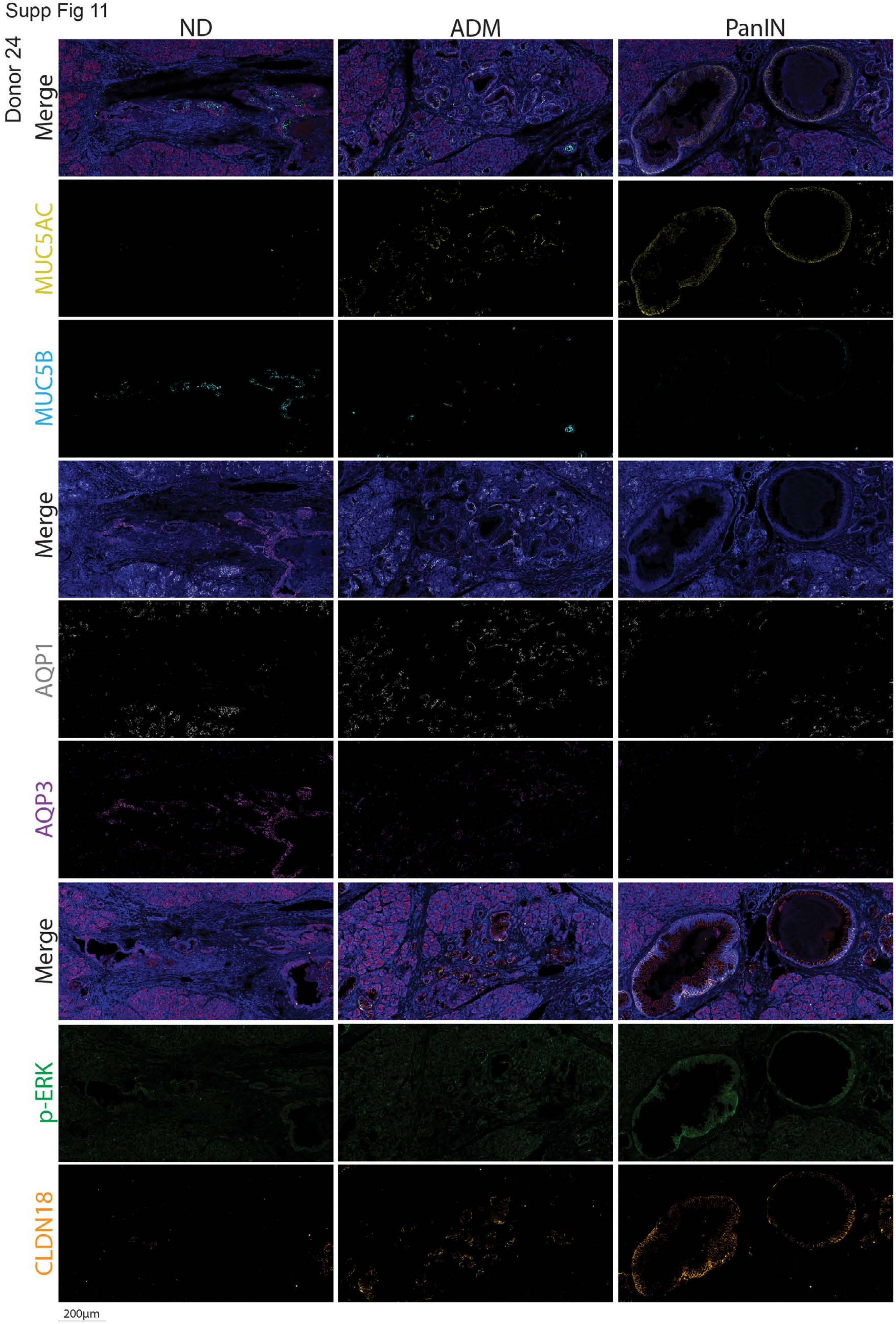

ND
ADM
PanIN
Donor 24
Merge
MUC5AC
MUC5B
Merge
AQP1
AQP3
Merge
p-ERK
CLDN18
200µm

Supp Fig 12

Donor 8

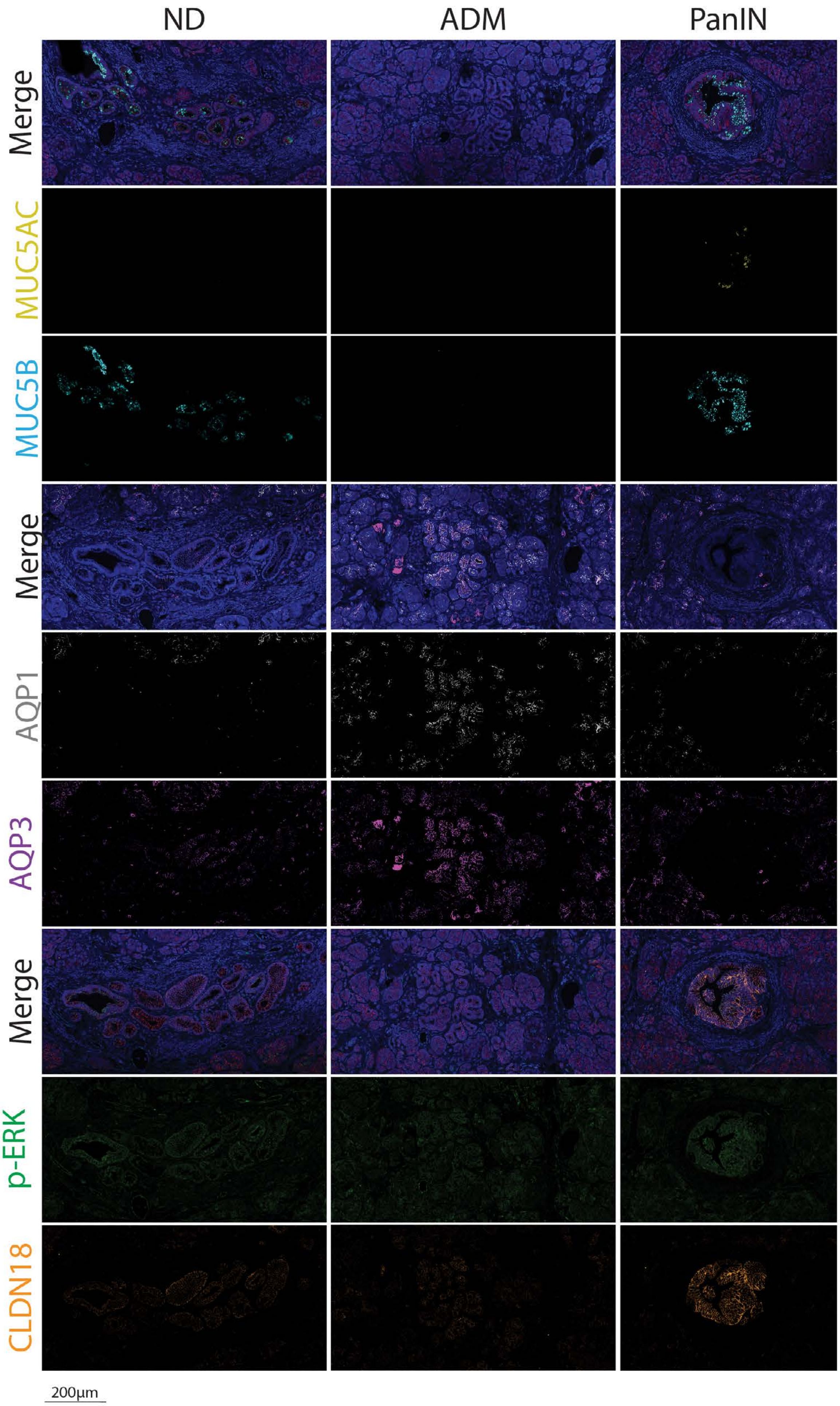

Supp Fig 13

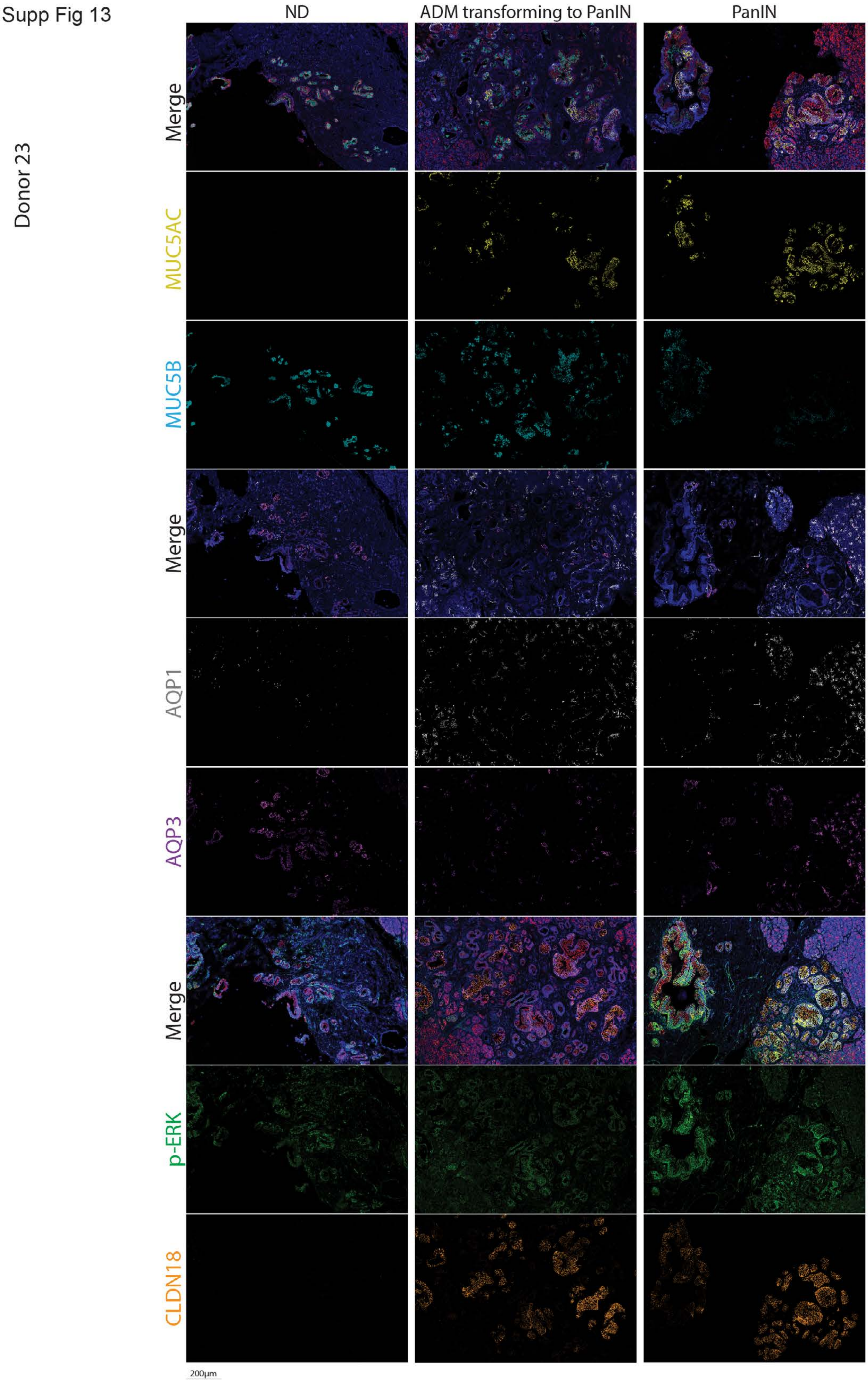
Donor 23
ND
ADM transforming to PanIN
PanIN
Merge
MUC5AC
MUC5B
Merge
AQP1
AQP3
Merge
p-ERK
CLDN18
200μm

Supp Fig 14

Donor 5

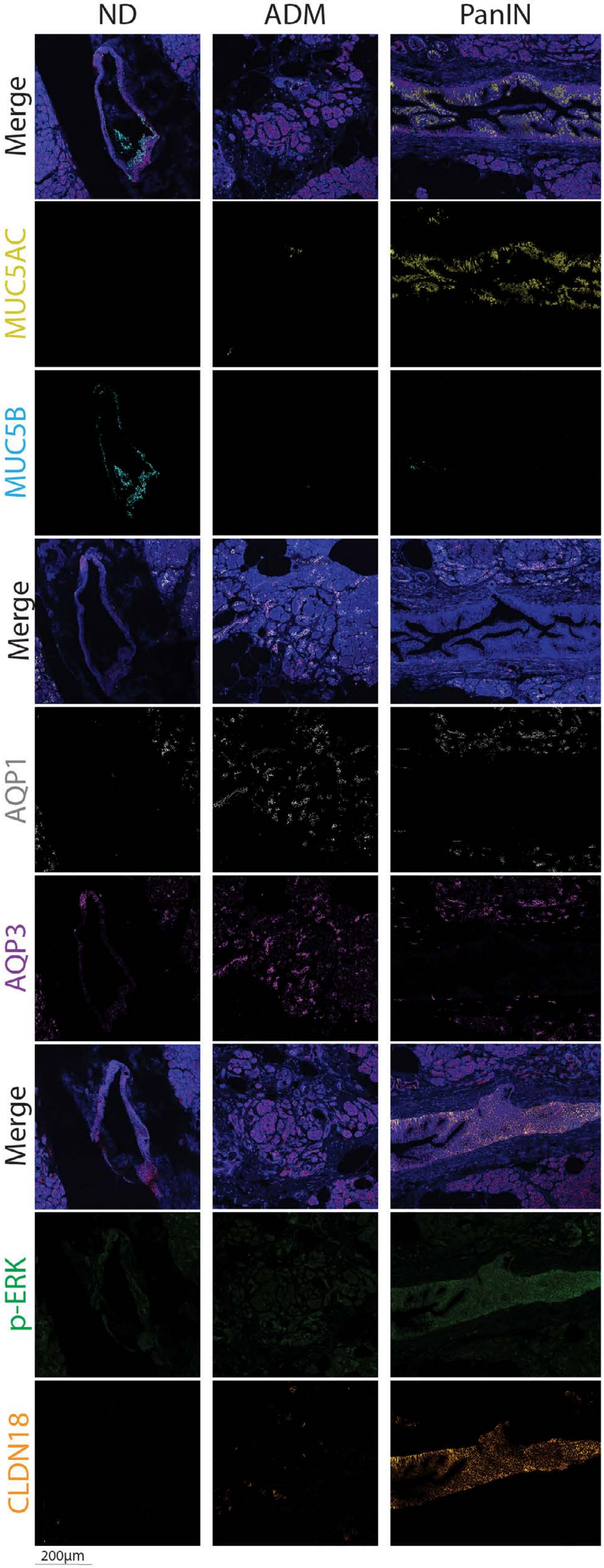
