## Supplementary Methods for "Analysis of donor pancreata defines the transcriptomic signature and microenvironment of early pre-neoplastic pancreatic lesions"

Immunofluorescence Assessment with Tyramide Signal Amplification

5µm thick formalin fixed paraffin embedded normal pancreas tissue sections were mounted on charged slides and baked at 60°C for 1 hour, deparaffinized and dehydrated. Slides were then washed with deionized water for 2 minutes followed by antigen retrieval at 96 °C with 10 mM sodium citrate (pH 6) for 8 min. Once the slides cooled down to room temperature, they were washed three times (2min per wash) with PBS and endogenous peroxides were quenched with 3% hydrogen peroxide for 15min. Then slides were washed three times with PBS. Tyramide signal amplification (TSA) kit (Invitrogen) was used according to manufacture’s instructions. Briefly, slides were incubated with blocking serum provided in the kit and stained with antigen-specific primary antibodies overnight at 4°C: Claudin 18 (1:100), Aqp1 (1:500), Aqp3 (1:200), Muc5ac (1:200), or Muc5b (1:200). The next day slides were washed three times with PBS and incubated in anti-rabbit secondary antibody for 1 hour. This was followed by an additional three washes with PBS. TSA conjugated to 488 or 555 fluorophores was applied for 10 minutes, followed by incubation with reaction stop solution for 5 minutes. Finally, slides were incubated with DAPI for 5 minutes and washed three times with PBS prior to being mounted with ProLong Diamond Antifade. High magnification images were obtained using confocal microscopy (Leica Stellaris 8). Antibodies used are listed in Supplemental Table 5.

*Single-cell RNA sequencing*

Samples were run using 50-cycle paired-end reads on the NovaSeq 6000 (Illumina) to a depth of 100,000 reads. Cell Ranger count version 6.0 was used with default settings, with an initial expected cell count of 10,000. The GCH38 refererence genome was used for aligment.Ambient RNA correction was done for each sample independently using SoupX(1). Briefly, SoupX estimates the ambient RNA profile from the cell-free RNA, then it estimates the cell-specific contamination fraction using a set of user-specified negative markers. Then, we used Seurat’s(2) recommended workflow for scRNASeq data processing and integration. Briefly, we removed low-quality cells with more than a 15% fraction of mitochondrial genes expression or less than 200 features (genes) detected. Expression was then log-normalized to the library size, the top 2000 highly variable features were extracted and scaled for downstream analysis. For samples integration, we used reverse principal component analysis (rPCA), as it is preferred for datasets with several mutually exclusive cell populations, and it scales better with a larger number of samples. Count matrices were integrated to account for batch effects. Using the integrated feature matrix, we ran PCA analysis to collapse gene expression to fewer principal components (35 principal components), then Louvain clustering was done to define different cell clusters. We used UMAP dimension reduction to create two-dimensional embedding for the cells. For cell-type annotation, we used a panel of previously known markers for different cell types in pancreatic tissue to define different cell populations. Differential cell-cell communication was performed using CellChat(3). Briefly, cell-cell communication was calculated between each cell-type pairs using the manually curated interaction database (CellChatDB) separately on tumor and healthy donor samples. Then differential cell-cell communication weight of tumor versus healthy was calculated for each cell-type pairs.

*Pseudobulk RNA Differential Gene Expression (DGE)*

We aggregated the counts from different samples for all (or a subset of) cells. We used DESeq2 for normalization and DGE analysis of the samples. We used PCAtools (<https://www.bioconductor.org/packages/release/bioc/html/PCAtools.html>) to perform and visualize PCA analysis on the samples and Pearson correlation to determine the correlation between the samples. Gene Ontology (GO) analysis was done using ClusterProfiler(4) package on the upregulated genes.

*Differential Abundance*

We used miloR to define differentially abundant cell populations (5). MiloR defines neighborhoods of cells using KNN graphs, after that, it uses negative-binomial regression to determine neighborhoods with differential abundance in one condition compared to the other. We set the K parameter for constructing the KNN graph to 60 as it yielded neighborhoods with a mean of 150 cells which is roughly equal 3*Number of samples as recommended by miloR authors.

*Gene set scoring*

For scoring different gene sets, we used AUCell(6) package. AUCell gives a score of a specific gene set for each cell using its raw counts, which makes it invariant to any downstream normalization or integration. Acinar, ductal, and tumor cell gene signatures were utilized from literature(7). We used MSigDB(8) Hallmark gene sets for KRAS activity score.

*Spatial Transcriptomics Data Analysis*

We used the GeoMxWorkflows package [https://bioconductor.org/packages/release/workflows/html/GeoMxWorkflows.html] for quality control and processing of Nanostring GeoMx data. Briefly, Quality control was performed to exclude the ROIs with less than 10% genes detection rate (less than 10% of the original WTA panel genes were detected) and exclude genes detected in less than 10% of the ROIs. Q3-normalization was performed for each ROI in the gene expression matrix followed by log normalization. For visualization purposes, we performed PCA analysis which clearly showed a batch effect. We used limma package(9) batch correction function to regress out the batch effect which showed clustering of ROIs from the same cell type together. To define cell-type-specific markers, we used a linear mixed model to perform DGE between each cell type and the rest of the ROIs, similar to how we define markers in single cells data. The top 20 significantly upregulated marker genes were then used as a geneset for scoring using AUCell(6). The scores are then scaled and visualized on the UMAPs of the acinar/ductal cells subset. To score different PDAC subtypes (Classical, basal, exocrine-like, ADEX) for each ROI, we used Gene Set Variation Analysis (GSVA)(10).
